# Computational and neural mechanisms underlying the influence of action affordances on value learning

**DOI:** 10.1101/2023.07.21.550102

**Authors:** Sanghyun Yi, John P. O’Doherty

## Abstract

When encountering a novel situation, an intelligent agent needs to find out which actions are most beneficial for interacting with that environment. One purported mechanism for narrowing down the scope of possible actions is the concept of action affordance. Here, we delve into the neuro-computational mechanisms accounting for how action affordance shapes value-based learning in a novel environment by utilizing a novel task alongside computational modeling of behavioral and fMRI data collected in humans. Our findings indicate that rather than simply exerting an initial or persistent bias on value-driven choices, action affordance is better conceived of as an independent system that concurrently guides action-selection alongside value-based decision-making. These two systems engage in a competitive process to determine final action selection, governed by a dynamic meta controller. We find that the pre-supplementary motor area and anterior cingulate cortex plays a central role in exerting meta-control over the two systems while the posterior parietal cortex integrates the predictions from these two controllers of what action to select, so that the action-selection process dynamically takes into account both the expected value and appropriateness of particular actions for a given scenario.

## 1 Introduction

In order to interact successfully with situations as they occur in the world, humans and other animals need to select particular actions from a very large set of possible actions based on which actions are most appropriate to the situation. One fundamental guiding principle for action selection, is that actions should be selected based on their expected value, that is by how much a particular action might increase an individual’s access to rewards, or decrease potential exposure to aversive outcomes [1–3]. A large literature has shed light on the neural and computational underpinnings of value-based action selection, including reinforcement-based mechanisms for learning which actions to select based on the future expected rewards they engender [4–6]. However, when encountering a stimulus for the first time in a particular context, the value of that stimulus is largely unknown and the brain needs to have a strategy to reduce the extremely large set of possible actions that could be selected to a tractable set of possible actions that could form the basis of subsequent trial and error learning. One putative mechanism for this is visual affordance [7–9]. Action affordances are features of stimuli in the world that suggest the appropriateness of particular actions, which can make the action selection problem more tractable. For example, a computer keyboard might suggest pressing or poking actions, a watering can might suggest a grabbing or clenching action, a pair of chopsticks might suggest a pinching action.

It has previously been found that affordance automatically potentiates particular actions that are compatible with properties an object or scene presented [8, 10–12]. Particularly, it has been suggested that the selection of visually guided actions is supported directly by affordance and that this affordance mechanism is in turn biased by other decision variables such as the value of choice options [13, 14]. However, the role that affordances might play in actually guiding learning during value-based decision-making is essentially unknown. A natural hypothesis is that affordance acts as an initial prior on value learning by, for example, subtly inflating the value of the afforded action during initial choice behavior so as to guide exploration, or alternatively, affordance might act as a constant yet moderate tug on action selection, persistently increasing the likelihood that an affordance-based action is selected in spite of the effects of value learning. Yet another possibility is that affordance-based choice and value-based choice operate as independent systems, competing for access to behavior. As we will see, our findings rather surprisingly support this latter possibility and suggest a role for a dynamic arbitration between independent expert systems implementing affordance and value-based choice.

To answer these questions we designed a novel behavioral task to probe the effects of affordance on decision-making. We adopted a computational model-driven approach – in which we specified a series of computational models to capture the different possible mechanisms by which affordance affects value learning, which we then systematically tested against human behavior. Further, we then measured brain activity with fMRI in order to identify the neural correlates of these putative computational processes. The overarching goal of the present study is thus to investigate how action affordances might potentially interact with value-based action-learning at behavioral, computational and neural levels.

In the behavioral task, participants were presented with pictures of an array of different visual objects. These objects were pre-selected to have specific action affordances based on the actions a separate group of participants rated to be most appropriate for a given object, out of three possible actions: pinch, poke or clench. In the task, when participants saw an object they could make one of those three actions in response, using a naturalistic (right) hand gesture (Fig. 1; Methods). We implemented a computer vision-based approach for the behavioral experiment, which classified a live video stream of the participant’s hand movements into one of the three gesture classes in real time (Fig. 1d; Methods). For a given object, one of these actions was associated with a higher probability of reward (winning money), while the other actions were associated with a lower probability of winning money. So for example, upon seeing a picture of a watering can, if the participant makes a poking gesture they would be most likely to win money, whereas if they make a pinching gesture they are less likely to win. Four objects (out of a set of 24 or 48 depending on the study), were encountered in a particular block of trials of 80 duration on average. Participants are instructed that their goal is for each object to select the action associated with the greatest amount of reward in order to obtain as many rewards as possible. Thus for each object, participants had to learn which action to select in order to maximize rewards. Crucially, for some objects, the most rewarded action was also the action that had the greatest affordance (congruent), while for other objects, the most rewarded action was not the action that had the greatest affordance (incongruent) (Fig. 1c). Furthermore, we included an orthogonal manipulation that controlled the reward probability such that half of the stimuli were associated with a high reward probability condition and the other half with a low reward probability condition, with the latter offering half the reward probability of the former (though in both conditions one of the actions available still had a higher reward probability than the other two). Through these manipulations, we could therefore assess the role that action affordance plays in guiding action selection alongside expected value, as well as characterizing how these two processes might interact. We ran two studies using this paradigm. An initial behavioral study (n=19) was followed by an fMRI study (n=30). Together these studies allowed us to investigate the behavioral effects of affordance on value-based learning and to then utilize computational modeling to gain insight into the computational mechanisms underpinning these interactions. Finally, our fMRI study allowed us to characterize the neural mechanisms implementing these computations.

**Fig. 1:**
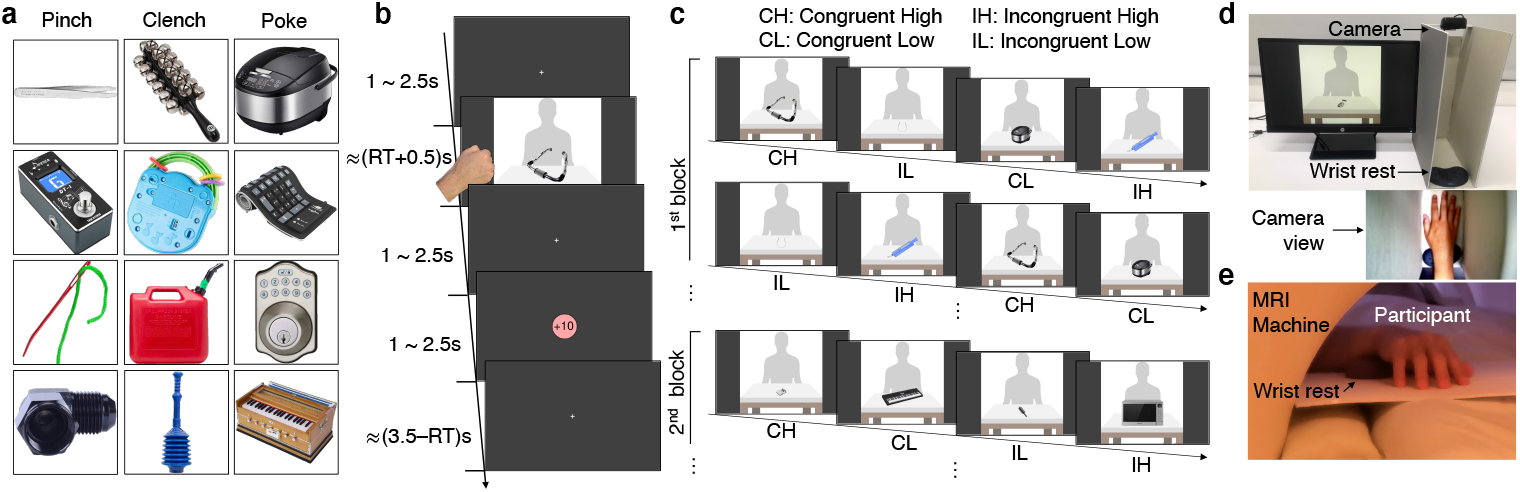
Decision-making task using naturalistic hand gestures and affordance-conferring stimuli. (a) Example stimuli associated with particular hand gesture affordances, determined from a separate online survey. (b) Example trial structure. A stimulus was displayed until an executed naturalistic hand gesture was registered. Each object was shown with a human silhouette background to give a naturalistic sense of the size of the object displayed. (c) Example block structure. Four objects were shown in each block and each object was associated with one of the 4 experimental conditions: congruent high (CH), incongruent high (IH), congruent low (CL), and incongruent low (IL) (see Methods for the details). (d) Experiment setting of the behavioral task. Participants made hand gestures inside the apparatus with a camera on top of it which sent a video stream to a server that classified hand movements into one of the three allowed hand gesture types in real time. (e) Implementation of the fMRI experiment. Participants lied down on the bed and mimed hand gestures on the plane which was placed on top of their bodies.

## 2 Results

### 2.1 Effects of affordance on reaction times and action selection

We first ascertained to what extent did the affordance properties of a stimulus influence response reaction times (RTs). We expected that choice of an action compatible with the dominant action affordance for an object would be associated with shorter RTs than choice of an affordance incompatible action regardless of the presence of ongoing value learning [10–12]. RTs were defined as the interval between the stimulus onset and initiation of movement, and were indeed found to be significantly shorter when making affordance-compatible actions compared to affordance-incompatible actions in both the behavioral and fMRI data even when participants were freely exploring the environment for finding the most rewarding hand gesture (Figs. 2a,f; for details on the reaction time measurement, see Methods). Moreover, the difference in RT remained consistent throughout. The average RT difference between the first 5 trials and the last 5 trials was not significantly different in either dataset (t(18)=-1.329, p=0.20 in the behavioral and t(29)=0.7592, p=0.45 in the fMRI data). We also found that the RTs were influenced by value-learning, in that participants were faster to respond to stimuli with higher expected value than stimuli with lower expected value (Supplementary Fig. 1). Thus, to disambiguate the effects of affordance on RTs from other possible confounding variables such as experimental condition, action type, choosing the most-rewarding action, and the progress of learning across trials, all of which are associated with value learning, we conducted mixed-effect linear regression which included each of these effects as potential confounding covariates. Even after accounting for these confounds, the effect of affordance on RT remained significant (Supplementary table 1; *β* = *−*0.026, *z* = *−*2.66, *p ≤* 0.01 for selecting affordance-compatible actions in the behavioral experiment, and *β* = *−*0.029, *z* = *−*3.02, *p ≤* 0.01 in the fMRI experiment, which indicates that RTs for affordance-compatible actions were about 2.7% shorter).

**Fig. 2:**
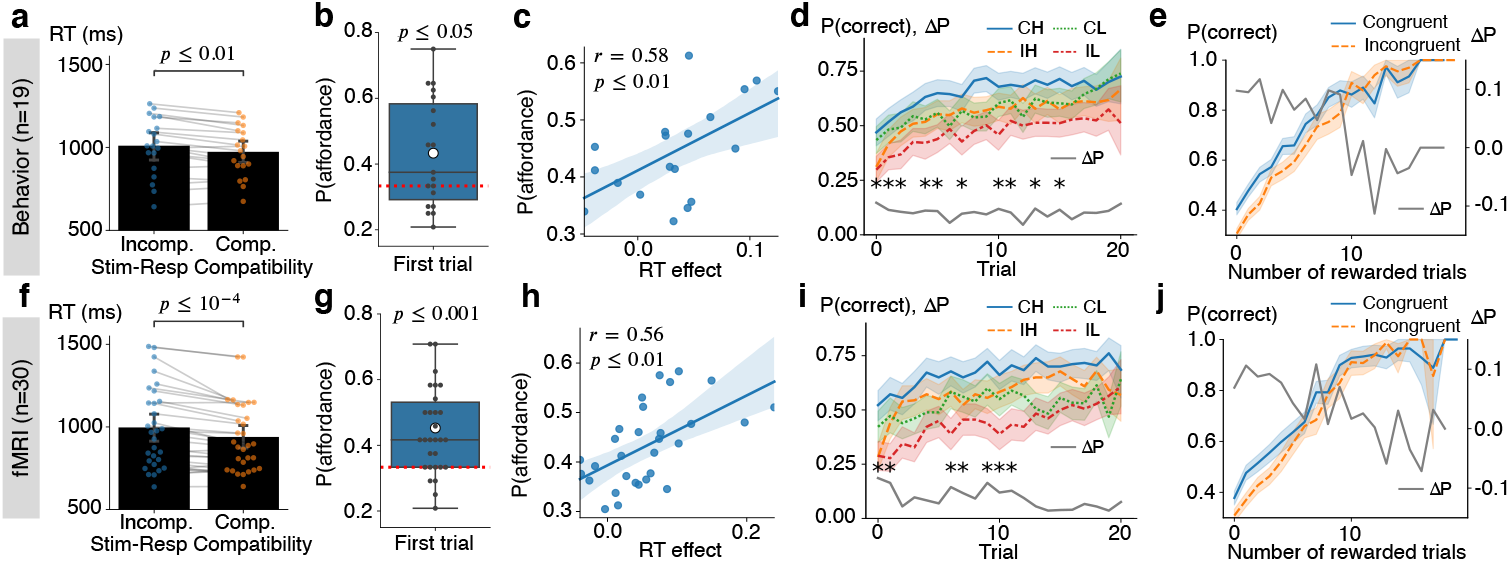
Behavioral effects of affordance during value learning and decision making. Upper-row plots show results from the behavioral study while the lower-row plots are from the fMRI study. (a and f). RTs were faster when the response was compatible with the affordance compared to when it was incompatible (paired-*t* tests; *t*(18) = 3.62 for the behavioral, *t*(29) = 4.68 for the fMRI experiment). RTs for each trial were averaged for each participant and trial types and the dots represent individual participants. (b an g) Initial responses to objects were significantly biased toward affordance-compatible actions. (*t* tests; *t*(18) = 2.62 for the behavioral, *t*(29) = 4.48 for the fMRI experiment). Dots represent the average frequencies for each participant and the red line is the chance level 1/3. (c and h) Correlations between the RT effect and the frequency of choosing affordance-compatible actions. (Pearson *r*) Dots represent each participant. (d and i) Learning curves averaged across blocks by experimental conditions. The star annotation presents the statistical significance of those difference between congruent and incongruent conditions in each trial (permutation tests; 5000 iterations each; *: *p <* 0.05 Bonferroni corrected) (e and j) Learning slopes averaged across blocks can be analyzed as a function of the number of rewarded trials. The gray lines in d,i,e and j show the choice accuracy difference between congruent and incongruent conditions. All the errorbar shows 95% interval of estimated statistics.

These results not only confirm that our experimental paradigm and stimulus set reliably induce affordance-related response preparation effects, but also reveal that the reaction time effect due to stimulus-response compatibility persists regardless of the effects of value-learning.

Next, we aimed to examine the effect of affordance on which action was chosen on a given trial. Specifically, we hypothesized that the affordance associated with a particular object would bias choices in favor of the afforded action, independently of the expected value of that action. To test for this, we analyzed the initial responses participants made to each object, as those are the actions not affected by value learning. The probability of selecting affordance-compatible actions as the initial response was significantly higher than chance (1/3) in both datasets (Figs. 2b and 2g). Furthermore, those initial actions were biased toward the afforded action that each object was selected to confer based on the initial affordance ratings we previously obtained in a separate sample (Supplementary table 2; *χ*^2^ test; null hypothesis was the probability distribution of choosing each action type independent of the affordance of objects calculated using the choice data; *χ*^2^(2, *N* = 304) = 9.33, *p ≤* 0.01 for pinch, *χ*^2^(2, *N* = 304) = 16.02, *p ≤* 0.001 for clench, *χ*^2^(2, *N* = 304) = 21.12, *p ≤* 10^*−*4^ for poke in the behavioral experiment; *χ*^2^(2, *N* = 240) = 17.91, *p ≤* 0.001 for pinch, *χ*^2^(2, *N* = 240) = 16.18, *p ≤* 0.001 for clench, *χ*^2^(2, *N* = 240) = 16.54, *p ≤* 0.001 for poke in the fMRI experiment). Moreover, the initial selection bias remained constant throughout the task for all new objects introduced over the course of the experiment (Supplementary Figs. 2a and 2c for details). This observation rules out an alternative explanation regarding the initial selection bias which posits that participants might have gradually inferred that the structure of the task is such that affordance-compatible actions are the most-rewarding action in half of the trials, and thus, the baseline probability for selecting affordance-compatible action would be 1/2 rather than 1/3. According to this hypothesis, the initial selection bias toward affordance-actions would be expected to increase as the task progresses as participants become more aware of the task structure with increasing experience. However, the actual data contradict this notion by showing that the initial selection bias remained stable and did not increase as the task progressed in both the behavioral and fMRI studies.

On top of that, we observed significant positive correlations across participants between these two distinct affordance-related bias effects on choice. We computed the degree of RT bias as the additional reaction time for executing affordance-incompatible actions relative to affordance-compatible actions 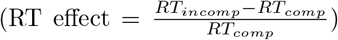 for each participant and compared it to the frequency of choosing affordance-compatible actions throughout the task. Significant correlations were observed between the two metrics in both datasets (Figs. 2c,h), and moreover, the RT effect positively correlated with the frequency of selecting the affordance-compatible action as the initial response to each object (Supplementary Figs. 2b,d; Pearson *r* = 0.41, *p* = 0.08 in the behavioral, Pearson *r* = 0.58, *p ≤* 0.001 in the fMRI experiment). These findings suggest that both forms of affordance-related biases on choice behavior are related and have a shared substrate.

### 2.2 Affordance influences value learning

The pronounced action selection bias toward the afforded action also influenced value-learning and contributed to the choice accuracy difference between congruent and incongruent conditions. To be specific, selection of the most rewarding action for each object was significantly greater in the congruent than in the incongruent conditions (*t*(18) = 2.50, *p ≤* 0.05 in the behavioral, *t*(29) = 3.25, *p ≤* 0.01 in the fMRI experiment). However, the bias toward selecting actions based on affordance was most evident in the early phase of the interaction with each specific object and diminished across subsequent trials involving that specific object. As demonstrated in Figs. 2d,i, statistical tests on the difference in choice accuracy between congruent and incongruent conditions in each trial revealed that the affordance effect was significant on early trials within a block but became less pronounced as learning progressed in both experiments.Because the exploration of choice options is biased by action affordance, participants are likely to need more trials to experience positive outcomes in incongruent than in congruent conditions. Therefore, the choice accuracy difference between congruent and incongruent conditions might be due to an affordance bias operating on the choice process rather than reflecting the effects of impaired value learning.

To analyze the influence of affordance on learning further, we examined the choice accuracy difference between congruent and incongruent conditions as a function of the number of rewarded trials previously encountered. The analysis indicates that learning slopes were actually steeper in the incongruent conditions, as evidenced by a decreasing pattern in the difference between the two learning slopes from each condition (Figs. 2e,j). Mixed-effect GLM analyses confirmed that the choice accuracy difference between the conditions is decreasing and that the incongruent condition has a steeper slope than the congruent condition (Fixed-effect coefficients for the gradient of the choice accuracy difference Δ*P*: *β* = *−*0.010, *z* = *−*3.16, *p ≤* 0.01 in the behavioral, *β* = *−*0.009, *z* = *−*2.97, *p ≤* 0.01 in the fMRI experiment; Supplementary table 3).

As previously mentioned, two possible explanations might account for this effect: first, it is possible that participants go through a longer exploration phase between receiving rewards in the incongruent conditions, which might support counterfactual learning during exploration, leading to steeper learning [15, 16]. In such scenarios, using a standard reinforcement learning (RL) model would suffice to account for the learning effect, as such a model could capture counterfactual learning by incorporating negative learning signals on the trials without rewards. Alternatively, it is plausible that behavioral adaptation based on reward history is more sensitive in the incongruent conditions. This could result from having a higher learning rate in incongruent conditions, or it could be due to the exertion of a higher level of cognitive control on incongruent conditions resulting in decisions that are more heavily reliant on learned values.[17–19]. As we will see, the computational modeling we implement supports the notion that cognitive control is allocated to balance the influence of affordance and value-learning in order to govern task performance (see below).

We also plotted the learning slope in high and low conditions as the function of the number of rewarded trials previously encountered. However mixed-effect GLM analyses revealed that the difference between high and low conditions does not decrease (Fixed-effect coefficients for the gradient of the choice accuracy difference Δ*P*: *β* = *−*0.006, *z* = *−*1.30, *p* = 0.20 in the behavioral, *β* = *−*0.010, *z* = *−*0.66, *p* = 0.51 in the fMRI experiment; Supplementary Fig. 3).

### 2.3 Dynamic meta-level control merging affordance-based and value-based decision-making best explains behavior

To investigate the underlying computational mechanism responsible for these behavioral effects, we implemented various computational models, fit those to participant’s behavioral data and then performed a formal model comparison (see Methods for details; Fig. 3a). We applied an RL model to capture value learning. We then modeled the degree of affordance assigned to each action for each object using the affordance-compatibility scores for the hand gestures for each object provided by the participants themselves after the completion of the value-based choice task.

**Fig. 3:**
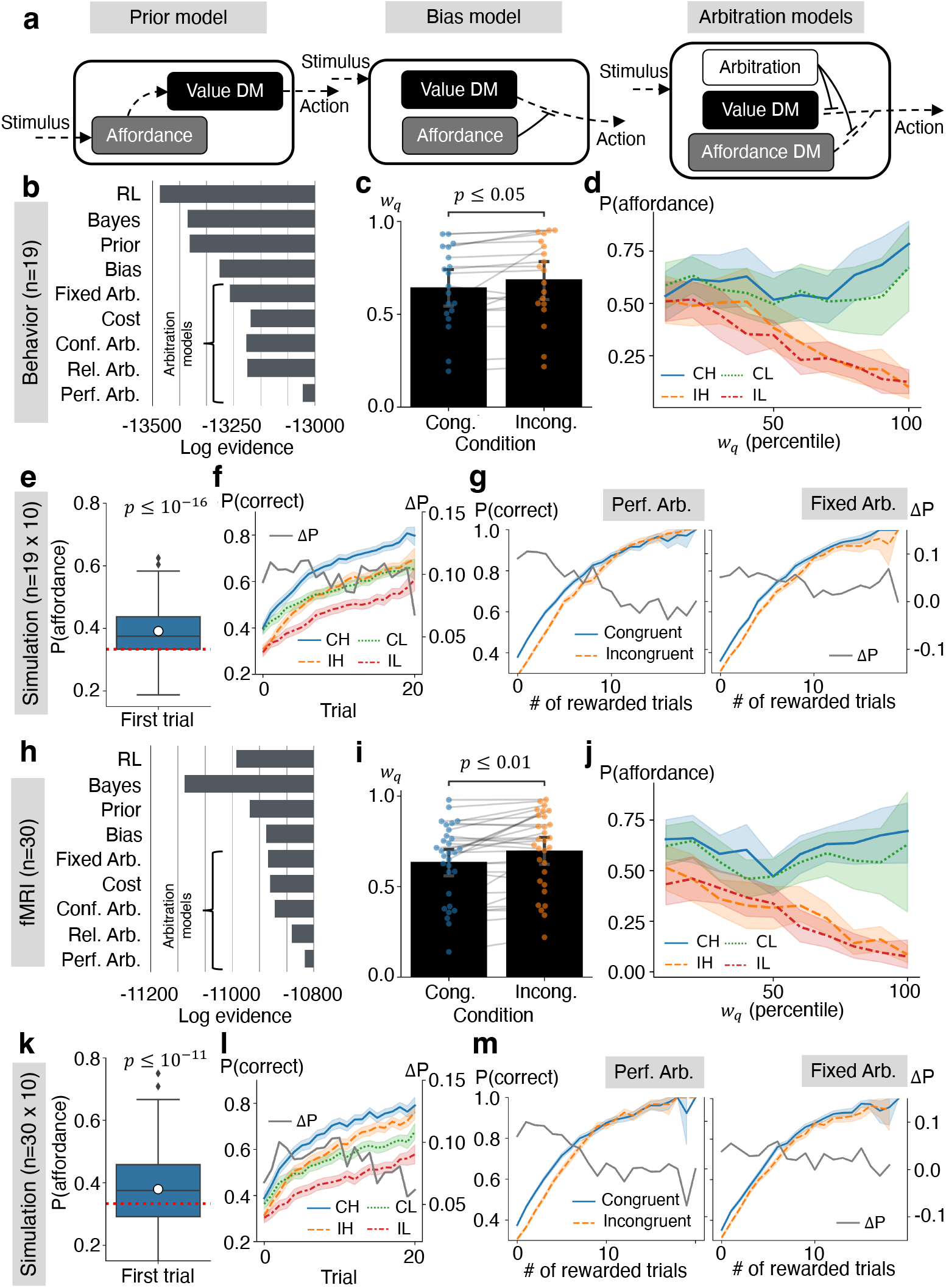
Computational model comparison, and the simulation results from the best model. Figs. 3b-g are from the behavioral study and their corresponding simulations while the Figs. 3h-m show the results from the fMRI study and their corresponding simulations. (a) Schematics of candidate computational mechanisms (Value DM: value-based decision-making; Affordance DM: affordance-based DM). See the main text and methods for details. (b and h) Log model evidence of the compared models. (c and i) Arbitration weights were affected by the affordance-value congruency so that the value-based decision-making was more favored in incongruent conditions. (paired-*t* tests; *t*(18) = 2.28 for the behavioral, *t*(29) = 3.59 for the fMRI experiment). Arbitration weights were calculated using the performance-based arbitration model for each trial and were averaged for each participant and condition. (d and j) Frequency of choosing affordance-compatible actions in the actual choice data is a decreasing function of the arbitration weight on the value-based decision-making. The arbitration weights were transformed into the percentiles within each participant. The tendency of choosing affordance-compatible actions more when the arbitration mechanism favors affordance-based decision-making was only evident in incongruent conditions as the responses based on affordance and value were indistinguishable in congruent conditions. (e and k) Simulated initial choice bias toward affordance-compatible action for each object. The performance-based arbitration models with individually estimated parameters were simulated 10 times each. (*t* tests; *t*(189) = 9.29 for the behavioral, *t*(299) = 7.21 for the fMRI experiment) (f and l) Simulated learning curves and their differences. (g and m) Simulated learning slopes and their differences as a function of the number of rewarded trials. All the error-bar shows 95% interval of estimated statistics.

One simple way in which affordance might influence choice is via a constant action-selection bias – essentially providing a constant push toward choosing the afforded action on each trial over and above other considerations such as value. Another possibility is that affordance acts as an initial prior operating on the initial value of the afforded action. If the afforded action is presumed to have a higher initial value than the other actions, this could produce a bias toward choosing that action more often at the beginning of a block. We implement both of these possible biases in separate RL models, either as a bias in the decision variable (bias model), or as a bias in the initial values assigned to an action before the onset of reinforcement-learning (prior model). We also tested the possibility that value learning was supported by Bayesian inference, rather than RL (Bayes model) [20].

In addition, based on the observation that the steeper learning slope in incongruent conditions (Figs. 2e,j) could be captured by cognitive control resulting in a greater focus on value in the incongruent conditions, we explored models inspired by the concept of a mixture of experts [21]. In this class of models, we assumed that two different decision-making systems are concurrently making predictions about the appropriate action for a given object. The first system is an affordance-based decision-making system, which simply makes choices in a manner proportional to the degree of affordance attached to particular actions, while the second system is a value-based decision-making system, by which actions get selected based on their learned expected values. Additionally, we assumed a meta-level controlling mechanism that arbitrates between the two systems and mediates the influence of the two systems in selecting actions. Four candidate arbitration mechanisms were tested.

The first model assumes that the outputs from the two systems are mixed with a fixed weight, which is conceptually similar to the bias model (Fixed arbitration model). We also tested a model that assumed affordance acts as a cost that makes the outcome from selecting affordance-incompatible actions less rewarding, thereby hindering selection of affordance-incompatible actions. This component was added on top of the fixed arbitration model (Cost model) [22]. The second type of arbitration model gave a boost in control to the value-based decision-making system, when the two systems made conflicting predictions, which mainly happened in incongruent conditions (Conflict-based arbitration model) [23–25]. The third model type assigned a greater degree of control to the system that had the lower level of prediction errors, or a higher level of reliability in its predictions, so that participants could minimize the uncertainty in predicting the values of each action. The reliabilities of each system were estimated using the absolute value of reward prediction errors (RPE) or affordance prediction errors (APE), which is the difference between the outcome and the affordance compatibility of the chosen action (Reliability-based arbitration model) [21, 26, 27].

The last type of model allowed for a larger influence from a system that had higher expected outcomes when the decision maker followed that particular system in making a decision [18, 22]. By doing so, the decision maker can maximize the outcome they can collect (see Methods for details). The performance of each system, or the expected outcome by following the specific decision-making system to a given object, was estimated using a method called inverse propensity scoring [28, 29], which was implemented in a form of delta rule supported by performance prediction errors (PPE) within each system. (Performance-based arbitration model; see Methods for details).

As illustrated in Figs. 3b,h, the performance-based arbitration model was found to best explain the actual choice data in terms of group-level log model evidence. Additionally, the Bayesian model selection results showed that the posterior model frequency and protected exceedance probabilities were in agreement with the above model comparison results (Supplementary Fig. 4). It is noteworthy that the task design used in this study had sufficient power to differentiate between the various decision-making models tested. For example, the performance-based arbitration model could be well recovered when the actual data generative process was based on itself. When the performance-based model provided the best fit for a given individual’s data, the probability of the performance-based model being the true generative model of the data was found to be 0.89 (Supplementary table 4; Methods).

In addition, the variables extracted from the best performing model suggest that the congruency between affordance and value determines how the meta-level controller weighs each system. Specifically, the arbitration weight on value-based decision-making was found to be higher in incongruent conditions (Figs. 3c,i; *t*(18) = 2.28, *p ≤* 0.05 in the behavioral; *t*(29) = 3.59, *p ≤* 0.01 in the fMRI data). We also observed that how often rewards were given on recent trials increased the arbitration weight toward value-based decision-making [22]. In the high conditions where rewards were given more frequently, the weight on the value-based decision-making system was higher (Supplementary Figs. 5a,f; *t*(18) = 2.53, *p ≤* 0.05 in the behavioral study; *t*(29) = 3.77, *p ≤* 0.01 in the fMRI study). Moreover, those trials with less weight on value-based decision-making were also trials in which the actual choice was compatible with the afforded action more often (Figs. 3d,j). We also observed that those participants who exhibited higher accuracy in selecting the most rewarding hand gestures were the individuals who also had a higher average arbitration weight assigned to the value-based decision-making system (Supplementary Figs. 5b,g; Pearson *r* = 0.62, *p ≤* 0.01 in the behavioral, Pearson *r* = 0.65, *p ≤* 0.001 in the fMRI experiment).

Furthermore, through model simulations utilizing the estimated parameters, we observed that the performance-based arbitration model could very closely replicate behavioral patterns found in the real choice data. In the simulated data, initial action selection was biased toward affordance-compatible actions and this bias was consistent across the blocks (Figs. 3e,k; Supplementary Fig. 7) as found in the real data described earlier. The simulated learning curves also exhibited a close correspondence with the actual learning curves in each condition, and the choice accuracy gap between congruent and incongruent conditions showed a tendency to be larger in the earlier trials of each object, which was consistent with the actual data (Figs. 3f,l; Fixed-effect coefficients for the gradient of the choice accuracy difference Δ*P*: *β* = *−*0.001, *z* = *−*2.26, *p ≤* 0.05 in the behavioral, *β* = *−*0.001, *z* = *−*1.50, *p* = 0.13 in the fMRI data simulation; Supplementary table 5).

It is notable that, apart from the performance-based arbitration model, simulations using other types of arbitration model did not show an initial action selection bias or a gradual decrease in the choice accuracy gap between congruent and incongruent conditions (Figs. 3e,f,k,l; Supplementary Fig. 8). Although the cost and the conflict-based arbitration models could exhibit several properties of the actual choice data with specific ranges of free parameters, simulations using the fitted parameters could not reproduce the patterns from actual choices (Supplementary Fig. 8). For example, the choice accuracy gap was increasing in the cost model, which is the opposite pattern from the real data. The conflict-based arbitration model did not show a bias toward affordance compatible actions in the initial trial. Moreover, the simulated choice accuracy in incongruent condition was even better or comparable in the conflict-based arbitration model which contradicts the actual data (*P* (*correct*|*incongruent*)*−P* (*correct*|*congrunet*) of the conflict-based arbitration model; *t*(189) = *−*3.12, *p ≤* 0.01 in the behavioral data simulation, *t*(299) = *−*0.69, *p* = 0.49 in the fMRI data simulation). The fixed and the reliability-based arbitration models could reproduce the initial action selection bias, but the gradient of the choice accuracy gap between congruent and incongruent conditions was marginal compared to that from the performance-based arbitration model (Supplementary table 5).

Additionally, when model-simulated learning curves were plotted as a function of the number of rewarded trials similar to Figs. 2e,j, the performance-based and reliability-based arbitration models could reproduce the real behavioral patterns, but not the fixed arbitration model (See Figs. 3g,m; Supplementary Fig. 9 and Supplementary table 6). For example, the difference between the congruent and incongruent conditions in terms of frequency of choosing correct actions as a function of the number of rewarded trials was only decreasing in the simulated data using the performance-based and the reliability-based arbitration models, but not in the fixed arbitration model’s simulated choices (Fixed-effect coefficients for the gradient of the choice accuracy difference Δ*P*: performance-based arbitration model’s *β* = *−*0.007, *z* = *−*6.64, *p ≤* 0.001 in the behavioral, *β* = *−*0.008, *z* = *−*8.81, *p ≤* 0.001 in the fMRI data simulation; reliability-based arbitration model’s *β* = *−*0.003, *z* = *−*3.08, *p ≤* 0.01 in the behavioral, *β* = *−*0.004, *z* = *−*4.28, *p ≤* 0.001 in the fMRI data simulation; fixed arbitration model’s *β* = *−*0.001, *z* = *−*0.91, *p* = 0.36 in the behavioral, *β* = 0.001, *z* = 0.62, *p* = 0.54 in the fMRI data simulation).

Nonetheless, the steeper learning slope we observed might be due to a potentially higher learning rate in the incongruent conditions. To explore this possibility, we fit the data with a fixed-arbitration model that incorporates two distinct learning rate parameters, one for incongruent conditions and another for congruent conditions. However, our analysis revealed no significant difference in learning rates between congruent and incongruent conditions (*t*(18) = *−*1.16, *p* = 0.26 in the behavioral, *t*(29) = 0.73, *p* = 0.47 in the fMRI data). Moreover, we could eliminate the possibility that the presence of counterfactual learning alone is sufficient to demonstrate the steeper learning slope, since all our tested models have such a property. Even with the implementation of counterfactual learning through the updating of values for unchosen actions in every trial, it is clear that counterfactual learning on its own falls short of replicating the actual behavioral data (see Methods and Supplementary Fig. 10). These results collectively indicate that the observed steeper learning slope in incongruent conditions is due to a dynamic arbitration mechanism that boosts the better working system for a given situation, rather than being a result of counterfactual learning in unrewarded trials or the effects of a larger learning rate in the incongruent conditions.

In addition, the non-decreasing difference between high and low conditions in terms of the frequency of choosing correct actions as a function of the number of rewarded trials could be captured by all tested arbitration models (Supplementary Fig. 11, Supplementary Table7 and Supplementary Note 2)

Consequently, through rigorous model comparisons and simulations, we identified performance-based arbitration as the best candidate model for explaining the behavioral data, both quantitatively and qualitatively. Next, we explored the neural implementation of affordance-based action-selection and its influence on value-based action-selection by utilizing computational variables from this model in the analysis of the fMRI data.

### 2.4 Neural correlates of affordance, value-learning, and action selection

We conducted a GLM analysis that included the chosen action’s affordance-compatibility score, action value and action selection probability from the performance-based model as parametric regressors to identify the regions associated with affordance and value-based decision-making respectively, as well as to uncover an action selection region responsible for integrating the predictions from these two systems to guide action-selection (GLM1; See Methods for details).

First of all, we found that the affordance-compatibility of the chosen action on each trial was encoded in the higher-level ventral visual stream such as V3 and V4 in the left occipital lobe which is the region that has been suggested to be responsible for affordance-perception or object recognition and processing physcial properties such as shape and size that are necessary for hand gesture control (Fig. 4a)[30–33] The chosen action value from the performance-based aribtration model was found in medial prefrontal cortex (mPFC) which is the region that has been implicated in the encoding of learned value (Fig. 4b) [34–36].

**Fig. 4:**
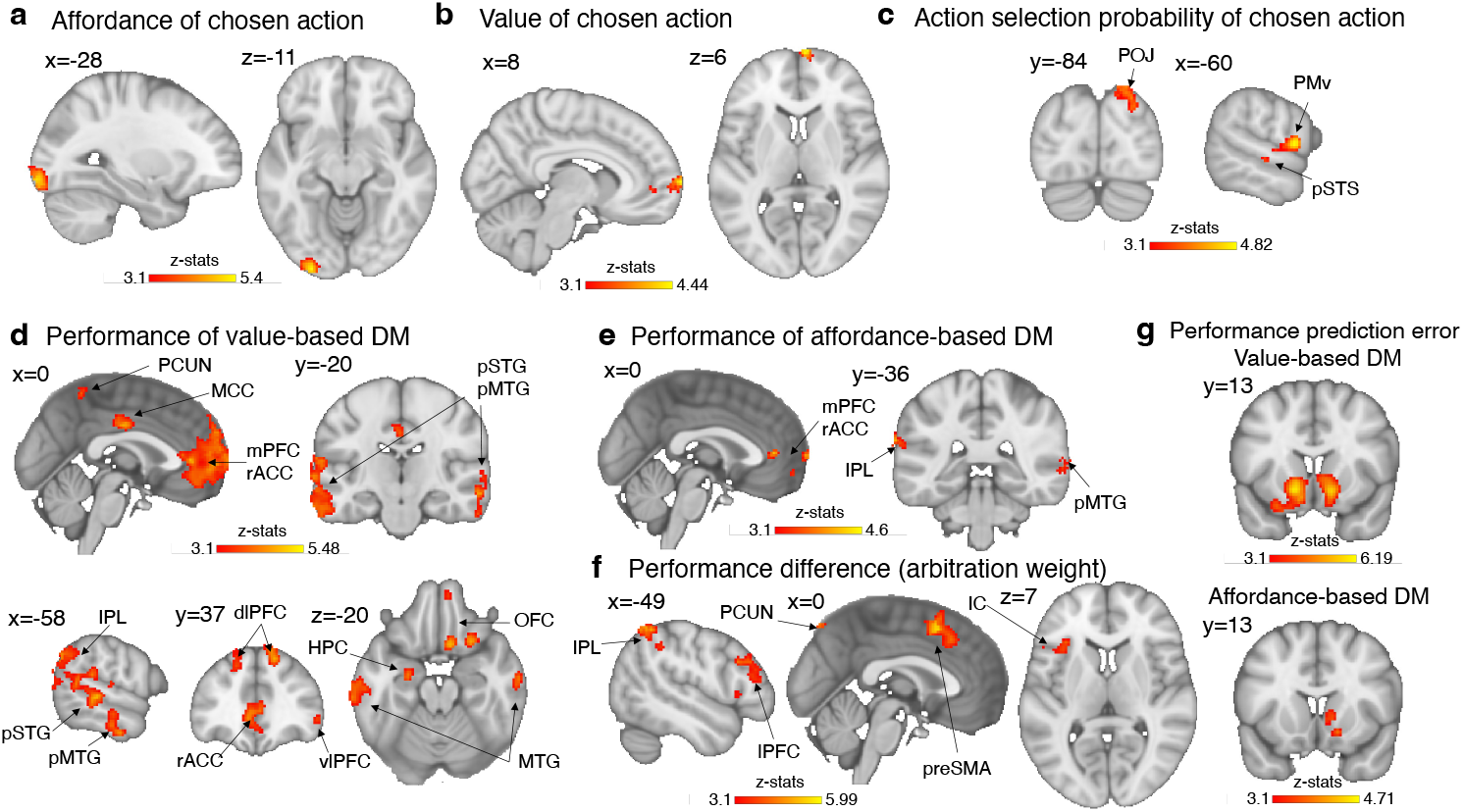
Neural implementation of performance-based arbitration. (a) Affordance-compatibility scores of the chosen action correlated with the high-level visual area including V3 and V4 in the left occipital lobe (b) The chosen action value was significantly identified in the mPFC. (c) Action selection probabilities of the chosen action were found in various regions of the cortical grasping circuit. (d) Performance of the value-based decision-making signals (e) BOLD signals of the performance of the affordance-based decision-making. (f) The fMRI correlates of the difference between the performances of two systems (*Perf*_*aff*_ *− Perf*_*q*_), which is directly related to the arbitration weight. (g) PPE signals for tracking performances of the two systems were identified in the striatum. All the results were cluster-corrected *p <* 0.05 with the cluster defining threshold *z* = 3.1.

Regions correlating with the action selection probability of the chosen action, which is the integration of the predictions from value-based and affordance-based decision-making systems were located in the posterior parietal cortex (PPC) near the parieto-occipital junction (POJ), ventral premotor cortex (PMv), and the posterior superior temporal sulcus (pSTS) all of which are parts of the cortical grasping circuit (Fig. 4c) [33, 37–39]. Notably, these identified regions are responsible for not only performing the hand gesture itself but also integrating different sensory modalities related to manual actions [40].

To ensure that regions correlating with the probability of the chosen action were not merely reflecting the effects of action execution vigor, we conducted an identical GLM analysis but incorporated RT as an additional parametric regressor [41]. These regions remained significant (at an uncorrected threshold of *p <* 0.001) suggesting that the regions are associated with the weighted sum of affordance-based and value-based decision-making systems rather than merely reflecting movement vigor (Supplementary Fig. 13).

Moreover, we extracted coordinates of the right-hand joints from the recorded hand videos and included them in the GLM as a parametric regressor in order to control for any potential confounding effects arising from the effects of hand movements per se (See Methods for details). While we found action selection related signals in various regions of the cortical grasping circuit, the actual motor implementation of the hand gesture was correlated with activity in the left primary motor cortex, left premotor cortex, left primary somatosensory cortex, and bilateral inferior parietal lobule (IPL) (Supplementary Figs. 14c-f).

Another GLM analysis using the second best fitting reliability-based arbitration model was conducted as well, which identified mostly overlapping regions for each of the cognitive variables (GLM2; See Supplementary Figs. 12a-c, and 13).

### 2.5 Neural implementation of the meta-level controller

Next, we probed for brain regions responsible for the meta-level computations involved in arbitrating between value and affordance-based decision-making systems. To achieve this, we utilized arbitration-related variables extracted from the performance model as parametric regressors in a GLM (GLM3; See Methods for details).

As illustrated in Figs. 4d,e, we found that the mPFC, rostral anterior cingulate cortex (rACC), posterior division of middle temporal gyrus (pMTG), and IPL to be associated with the performances of both systems. The posterior division of superior temporal gyrus (pSTG), dorso and ventro lateral prefrontal cortex (dlPFC, vlPFC), mid-cingulate cortex (MCC), orbitofrontal cortex (OFC), precuneous cortex (PCUN), and hippocampus (HPC) were correlated with the performance of value-based decision-making but not with the performance of affordance-based decision-making.

In addition, we observed that the signal corresponding to the difference in performance between the two decision-making systems (*Perf*_*aff*_ *−Perf*_*q*_), which determines the arbitration weight, was present in the pre-supplementary motor area (preSMA), lateral prefrontal cortex (lPFC), and insular cortex (IC) (Fig. 4f). We also found that the IPL and PCUN were regions correlated with the difference in performance between the affordance and value-based decision-making systems. These regions have been shown to be associated with the implementation of cognitive control [18, 42–45]. Interestingly, our findings indicate that the regions identified were less activated when the value-based decision-making system was allocated more weight.

Similar to the RPE signals (Supplementary Fig. 14a), the variables responsible for updating arbitration weights, which are also outcome-dependent prediction error signals, were found in the striatum (Fig. 4g). Specifically, the PPE of the value-based decision-making system was prominently encoded in the ventral striatum, while the PPE of the affordance-based decision-making system was found in the dorsal striatum.

In addition to investigating neural correlates for the performance-based arbitration model, we also tested for regions correlating with reliability-based arbitration which was the second best fitting model in our model comparison (GLM4). We also found clear neural correlates of reliability-based arbitration signals (Supplementary Figs. 12d-g and Supplementary Fig. 14b). In particular, we found the regions that correlate with the difference between the reliabilities of the two systems largely coincide with those identified using the performance-based arbitration model, particularly the preSMA. It is note worthy that the average correlation between the arbitration variables of the two models were small(*r* = 0.196 in the fMRI data and *r* = 0.159 in the behavioral data). Therefore, the most plausible explanation is that each model captures distinct variance in activity in the preSMA. These findings indicate that the brain keeps track not only of the performance of the different systems but also of their reliability, suggesting that both variables might ultimately be taken into account during the arbitration process.

### 2.6 Better task performance is linked to a more robust representation of arbitration variables in meta-level controller regions

Next we explored the relationship between individual differences in task performance and representation of the arbitration variables. We found that individuals with higher task performance demonstrated more robust representations of the arbitration variables such as the performances of the two systems and the difference in performance between the two systems in brain regions involved in encoding arbitration-related variables. To be specific, we found positive correlations between each participant’s propensity to choose the most rewarding actions and the extent to which BOLD activity in the identified arbitration regions can be explained by our model-derived arbitration variables.

For instance, in those participants who tend to be more accurate in choosing the correct actions, the computational variable corresponding to the difference in performance between the two decision-making systems provided a better account of BOLD activity in the arbitration regions shown in Fig. 4f (*r* = 0.47, *p ≤* 0.01; Fig. 5a, See Methods for details). Moreover, in those participants who achieved higher accuracy in choosing the most rewarding actions, brain regions involved in representing the computational variables corresponding to the performance of each individual system were more correlated with the model-estimated performance variables (*r* = 0.38, *p ≤* 0.05 for the performance of the affordance-based system; *r* = 0.43, *p ≤* 0.05 for the performance of the value-based system; Fig. 5a). However, the strength of the neural representation of the *within* system computational variables produced by each system relevant for decision-making did not correlate with the participants’ accuracy (*r* = 0.23, *p* = 0.22 for the affordance; *r* = 0.08, *p* = 0.68 for the value; Fig. 5a). These findings, suggest that more robust neural representations of performance-based arbitration are associated with better behavioral performance on the task, providing additional evidence in support of the role for an arbitration process in mediating effective interactions between value-based and affordance-based action selection (Analogous results were also found for the reliability-based arbitration model; see Supplementary Fig. 15).

**Fig. 5:**
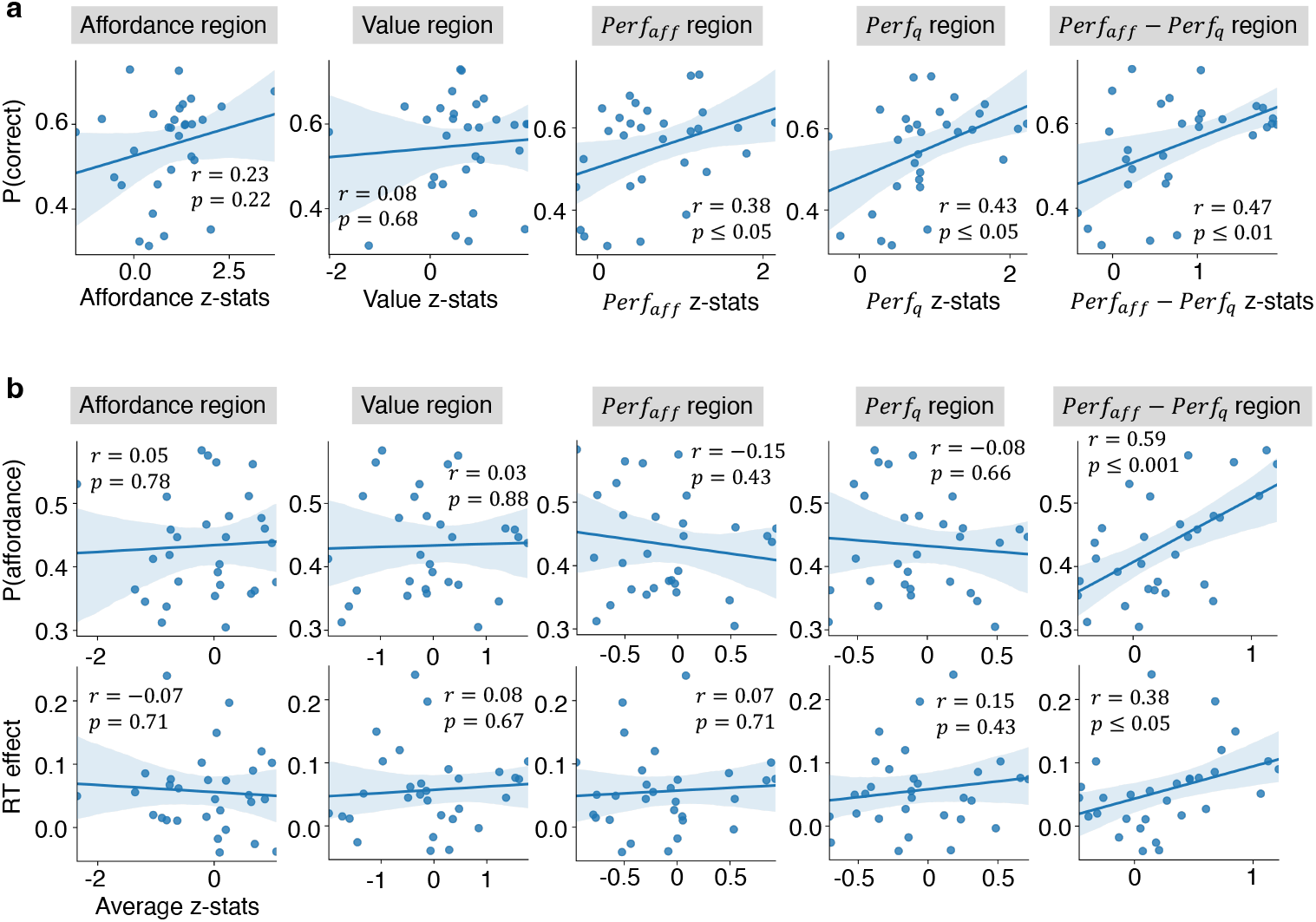
Correlations between behavioral effects and neural representations. (a) Correlations between the tendency to select the most rewarding actions and the strength of the neural representation of cognitive variables by functionally defined regions of interest. (b) Correlation between the affordance effect on the behavior and the increased BOLD activity when executing affordance-incompatible action by functionally defined regions of interest. The x-axis is the average z-statistics of the contrast between the affordance-incompatible and affordance-compaitble action execution from the GLM5 across the voxels within the ROIs. RT 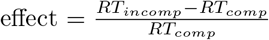. Each dot represents an individual participant. See Methods for the details.

We also found that those individuals with a stronger behavioral affordance effect exhibited increased activation of performance-based arbitration regions when executing affordance-incompatible actions. Such individuals likely have to exert stronger behavioral control to suppress the prepotent affordance compatible action compared to individuals with a lower overall affordance effect in their behavior. To test for this, we looked at the relationship between the overall proportion of affordance compatible actions chosen by each individual and activation on trials where the affordance incompatible action was chosen compared to when it was not chosen (GLM5; See Methods for details). A significant correlation was observed in performance-based arbitration regions (*Perf*_*aff*_ *− Perf*_*q*_ regions in Fig. 4f; *r* = 0.59, *p ≤* 0.001; Fig. 5b). A similar effect was found when examining the relationship between the reaction time increase when executing affordance-incompatible actions compared to affordance compatible reaction times, and activity in those areas (*r* = 0.38, *p ≤* 0.05; Fig. 5b; See Methods for details). These findings suggest that regions involved in arbitration including the preSMA, have an important role in suppressing the prepotent affordance response, especially in those individuals who have a stronger overall bias toward affordance effects [42].

## 3 Discussion

We provide evidence that action affordance plays a key role in shaping value-based action-selection and learning in humans. Action affordance and value-based decision-making were found to interact to guide behavior. Affordances were found to both influence reaction times, such that actions compatible with the affordance are selected more rapidly regardless of the effects of ongoing value learning, as well as biasing choice toward the afforded action, an effect that was most prominent in the early trials.

In order to understand the mechanism by which action affordance influences instrumental value learning, we implemented and tested a series of computational models. We found that the effects of affordance on choice behavior is not well described by either a simple bias in action-selection or an initial prior boosting the value of afforded actions. Instead, a form of dynamic arbitration was found to best explain behavior on the task. According to this framework, two separate systems operate in parallel, a value-based system and an affordance-based system, and the determination of which system contributes most to behavior at any one time is based on estimates of the relative performance of the two systems. This neuro-computational mechanism provides a potential explanation for the problem of how human learns appropriate actions in the absence of clear information about their value. Given that a very large family of actions could be implemented in any given situation, it is highly beneficial for the set of actions available for selection to be constrained by properties of the visual environment such as object affordance. This can help make the action-selection problem more tractable, and provide a scaffold for selecting actions so as to explore and learn about value during initial exploration.

In the brain, consistent with our computational model-based findings, we found evidence for the existence of distinct affordance-based and value-based decision-making systems. While the value of the chosen action was found in medial prefrontal cortex, the affordance of the action chosen on a given trial showed a positive correlation with activity in the visual pathways of the occipital cortex, including regions V3 and V4 [30, 34–36]. This finding suggests that the occipital visual areas, crucial for recognizing an object’s physical properties and identity, are more engaged when an action compatible with the object’s affordance is chosen [31–33]. Conversely, when an action irrelevant to the object is selected, these characteristics are less involved in the final action selection and motor execution process.

Moreover, our study revealed that the overall integrated choice probability, which is the integration of value and the compatibility of certain actions to the shown object, is associated with activity in the PPC. The PPC is known for representing object-associated hand movements and has been identified as a key area in reward-based decision making, especially concerning action values[35, 46–51]. Thus, our finding aligns with previous findings and supports the perspective that the PPC plays a critical role in integrating multimodal information necessary for movement execution [52]. It also unifies two different literatures on visually guided motor control and economic decision making, emphasizing the PPC’s comprehensive function in these processes.

In addition to identifying regions correlating with each strategy and their integration, we also looked for evidence of brain regions involved in meta-control over the two underlying action-selection systems. Specifically, we identified a network of brain regions tracking and comparing the performance of both strategies, including the anterior cingulate, pre-SMA, lateral prefrontal cortex, and inferior parietal lobule, regions that have been found to be involved in cognitive control in a number of previous studies [18, 42–45]. Moreover, activity within the arbitration regions served as an indicator of the likelihood that participants would choose affordance-compatible actions, as well as the extent of the stimulus-response compatibility effect on reaction times. Additionally, we found that the more activity in regions representing arbitration-related variables correlated with the relevant arbitration variables across participants, the more optimal a participant’s behavior was in terms of selection of the most rewarding actions.

This finding therefore suggest a crucial role for the arbitration process between affordance and value-based decision-making in guiding optimal behavioral outcomes. The finding of a role for an arbitration scheme governing the contribution of value-based and affordance-based choices to behavior is consistent with a mixture of experts framework [21]. In the present study, we found evidence for a role of performance-based arbitration over and above reliability-based arbitration, which we have previously investigated in relation to its role in allocating control between other strategies such as model-based and model-free RL and different forms of observational learning [26, 27]. While these two arbitration concepts are closely related as they are both concerned with how well a particular expert system is doing in making predictions, the precise computational variables underpinning the arbitration process is an important research question. In the present study, reliability-based arbitration was the second best performing model, outperformed only by the performance-based arbitration model. Moreover, we found evidence for reliability-based arbitration signals in the brain alongside performance-based arbitration signals and those two arbitration signals capture distinct variance in the activity of the identified arbitration regions which includes the preSMA. It is possible that both performance and reliability are playing a role in guiding arbitration between these different expert systems.

It is important to note that the arbitration process described here can be seen as a form of cognitive control. Central to our model is the concept of directing control towards the system that promises a higher expected return. This idea aligns with the Expected Value of Control (EVC) model which allocates cognitive control based on the control’s effectiveness such as expected value and cost [22]. According to the model, the demand for control and deciding the control intensity can be evaluated by tracking the conflict between task relevant and irrelevant information as well [24]. However, our findings suggest that a conflict-based arbitration model, rooted in conflict monitoring theory, does not most effectively explain choice behavior in value learning contexts, despite the proposed significance of conflict signals in control allocation. In contrast, our model emphasizes allocating control towards the system with a higher expected return, a concept resonating with the EVC model’s focus on control effectiveness.

Furthermore, the well-established link between perception and action extends beyond the concept of affordance. The automatic potentiation of likely actions has also been explored from an information processing viewpoint, indicating that motor and visual representations share a common representational space [53, 54], and from attention theory, which suggests that attention to a spatial location primes corresponding action plans toward that location [55, 56]. While efforts have been made to differentiate the affordance effect from the Simon effect [57–59] the necessity of affordance in explaining stimulus-response compatibility is still debated [60]. Notably, the affordance effect is significant primarily when motor representation, aligned with an object’s affordance, is triggered by movement intention rather than solely by the visual stimulus [61, 62]. Our experiment which required movement intention for completing each trial, is particularly relevant in this context. Recent studies also suggest that affordance, or plausible actions triggered by a stimulus, impact attention allocation and other cognitive processes such as working memory and visual perception. This indicates that explaining affordance through spatial attention alone is insufficient, and that a bidirectional process must be considered [63–65]. Our computational approach was not focused on the mechanisms behind affordance-induced stimulus-response compatibility but instead on how automatically activated actions influence learning.

Another pertinent question concerns how affordance-based action-selection relates to other forms of action-selection that are thought to exist alongside value-based decision-making. The most obvious comparison is with habits. Habits are suggested to be formed via repeated reinforcement of stimulus-response associations [66]. Here, the affordances associated with particular objects were not acquired in the experiment through trial and error reinforcement, because they were manifested on the very first trial that each object was encountered and were by design kept orthogonal to the reward contingencies. Thus, they are not “habits” in the traditional sense as typically studied in the lab. However, it is possible that affordances do correspond to a type of stimulus-response association that has been historically acquired over the course of development as individuals interact with objects in their environment and learn to implement specific physical actions in response to them. In that sense, these affordances could be thought of as very well-learned habits. However, our fMRI study revealed evidence for the neural implementation of affordances and their influence on action-selection in occipital and parietal cortices, and not in areas traditionally associated with habitual action-selection such as the posterior putamen [67, 68]. It is possible that such extremely well learned habits come to eventually depend on the cortex and not the basal ganglia. However, within the dorsal striatum, we did find learning update signals related to meta-control that steers individuals to favor affordance-based decisions – suggesting that the basal ganglia may play a role in updating control signals related to the arbitration process between strategies, over and above its contributions to implementing individual strategies.

Alternatively, affordance-based influences on action-selection could be implemented via a much more dynamic visuomotor computation, in which specific visual features of an object guide on-line action-selection computations in which the most relevant actions for interacting with a particular object are decided upon and implemented [69]. Further investigation of the specific neural computations unfolding during affordance-based action selection could help discriminate the underlying mechanisms.

We acknowledge a potential limitation in our study arising from the use of object images instead of real objects. Images might primarily convey the object’s semantics or ‘stable affordance’, constructed from people’s familiarity with the object, and not fully represent the physical properties like the object’s orientation and proximity to participants, which are also important factors in affordance perception [70]. Additionally, there is a possibility that the different neural circuitry has been involved in the current study compared to using the real object. For example, researches have shown that interacting with real objects and images involve distinct visuo-motor circuitry and object pictures are rather processed conceptually or as the object word [71–73]. Furthermore, fMRI studies have identified parieto-occipital cortex regions sensitive to the physical properties of real objects [74–78].

Nevertheless, our data suggest that the effects we found about hand gestures associated with objects was influenced not just by participants’ familiarity and the semantics of objects, but also by their physical properties inferred from visual features of the displayed objects. The familiarity scores participant reported about stimuli used in the behavioral and fMRI tasks varied significantly (66.45 *±* 24.62 on a 0 to 100 scale, Supplementary Fig. 16) and the difference in probabilities of choosing affordance-compatible actions in the initial trial upon seeing unfamiliar objects (familiarity score *<* 66.45) and familiar objects (familiarity score *≥* 66.45) was not significant (t(18)=1.529, p=0.144 in the behavioral, t(29)=0.001, p=0.996 in the fMRI), despite a general bias towards the affordance-compatible actions in the first trials. This indicates that the choices were influenced by their physical properties discernible in the images, beyond just familiarity or object identity. Furthermore, the stimuli were not arbitrarily selected but annotated by online participants viewing the same images as the on-site participants. From approximately 1000 stimuli, 48 objects significantly associated with specific hand gestures were chosen for our experiments. Thus, the affordance labels used in our study can be considered stable, taking into account the object’s orientation and virtual proximity as displayed in the stimuli.

Additionally, the automatic potentiation of action and its behavioral and neural effects have been observed even with photographic representations of objects [11, 79]. While planar presentation may affect visuo-motor processing, stable affordance aspects such as mechanical and functional knowledge about an object are likely unaffected [80]. Consequently, our findings remain valid within the context of studying the effect of stable affordance, predominantly derived from object semantics, in value learning. However, future research utilizing real objects will be helpful for studying affordance in more ecologically valid and realistic settings.

In conclusion, the present study provides evidence that human value learning is guided not only by the value of particular actions, but also by the visual affordance of objects. Rather than affordance acting as a simple bias in decision-making or prior in value learning, instead we find that affordance and value-based decision-making are best viewed as distinct expert systems that interact by means of a determination about which system is performing best in obtaining rewards.

## 4 Methods

### Participants

We recruited 21 and 32 healthy participants for the behavioral-only experiment and the fMRI experiment respectively. 2 participants were excluded from the behavioral and another 2 from the fMRI experiment because they didn’t complete the study. Therefore, data from the remaining 19 (8 females, 18 right-handed; 1 ambidextrous, 18 *∼* 24 years: 6; 25 *∼* 34 years: 10; 35 *∼* 44 years: 1; 45 years or above: 2) and 30 (13 females, 28 right-handed; 2 ambidextrous, 18 *∼* 24 years: 12; 25 *∼* 34 years: 12; 35 *∼* 44 years: 4; 45 years or above: 2) were used for the analyses. Before taking part in the experiment, all participants were assessed to ensure that those with the history of neurological or psychiatric illness were excluded. All participants provided their informed consent, and the study was approved by the Institutional Review Board of California Institute of Technology.

### Stimuli

To implement the paradigm, we began by creating a set of stimuli. The stimulus set consisted of various object images, each of which was designed to engender a specific hand configuration among four hand movement classes known to be employed during object interaction: pinch, clench, poke, and palm (e.g., a button-shaped object that affords poking)[81]. We first obtained 1000 images of around 900 unique objects and built a set of visual stimuli by superimposing those object images onto a human silhouette image. We designed the stimuli in this way to ensure that the visual stimuli retain information about the object’s size.

Then we used those edited photographs in an online task on Amazon Mechanical Turk (M-Turk). Each image was displayed 4 times during the task to collect annotations on the suitability, or affordance-compatibility score, of 4 hand movements to the object displayed. Within a trial, the image with the human silhouette was displayed for 2 seconds and the object’s zoomed-in image was shown for one second along with a text of one of the 4 hand gestures. Then while the zoomed-in image was on the screen for additional 8 seconds, M-Turk participants responded how suitable it is to pinch, clench, poke or palm the object shown by sliding a bar ranging from “Very unsuitable” to “Very suitable”. Additionally, we asked about their familiarity with the object 4 times. The affordance-compatibility and familiarity scores were both converted to 0 to 100 scale later. Throughout the task, 4 catch questions were used to check the participants’ attention.

We recruited 227 participants online, but only 160 of them were included in the data analysis as we excluded those that didn’t pass at least 2 catch trials. Each of the participants was asked to annotate 50 objects. On average, each object was annotated by 7.2 individuals.

Then we chose objects based on their 0-100 affordance-compatibility scores. For example, if an objects’ average affordance-compatibility score of pinch was greater than 50, and the mean score of pinch was significantly larger than the mean scores of other actions (independent t-tests, *p <* 0.05 uncorrected), the object was chosen as a pinch-affording object. By doing so we obtained 21 pinch-, 102 clench- and 16 poke-compatible objects. Then we randomly selected 16 items from each set, yielding 48 stimuli for the main task (Fig. 1a).

### Task

We designed a variant of a 3-armed bandit task in which the available actions are natural pinch, clench, or poke gestures made by the right hand, which yields binary outcomes. We used 48 visual object stimuli for the behavioral experiment and 24 object stimuli (a subset of the 48 stimuli) for the fMRI experiment. On each trial, one of the stimuli showing an object with a human silhouette was displayed, and a participant mimed one of the three hand gestures while maintaining their wrist position on the provided wrist pad (Figs. 1a,e). The images used here (including the human silhouettes) were identical in form to the selected sub-set of images initially presented in the online survey. Participants were provided with instructional videos showcasing example hand movements corresponding to each gesture type to ensure that they had a clear understanding of the hand gestures under consideration. Participants were instructed that the reward probability of each hand motion is different for each object, and that they need to figure out the most rewarding hand gesture for each object by trying different hand motions. Each category of hand postures corresponded to actions with different reward probabilities, and an object was shown in a trial to evoke one of the three hand gesture affordances. There were 4 experimental conditions in this paradigm: congruent high, congruent low, incongruent high, and incongruent low conditions. In a congruent trial, an action that was consistent with the presented stimulus’ affordance had a high reward probability, whereas in an incongruent trial, the opposite applied. Each hand position had a reward probability of 0.8, 0.2, or 0.2 in the high condition and 0.4, 0.1, or 0.1 in the low condition.

The task consisted of a 2-day experiment for the behavioral-only study and a 1-day experiment for the fMRI study, with 76*∼*84 trials (on average 80 trials) per block, and 6 blocks per day, totaling 960 trials for the behavioral study and 480 trials for the fMRI study. In each block, 4 distinct objects were displayed using the event-related design. Each object corresponded to one of the 4 conditions and was shown for an average of 20 trials (ranging from 19 to 21 trials). All objects were displayed once in every 4 trials in random order. Everyone in the behavioral-only and fMRI experiment was shown the same set of 48 or 24 objects respectively, but the associations between experimental conditions and stimuli was randomized across participants. For the counterbalancing, at most two objects within a block had the same affordance and at most two objects within a block had the same most-rewarding response. Also, the two congruent condition objects within a block had different affordances and the two incongruent condition objects had different affordances and different correct responses from each other.

Each trial lasted between 7 to 11.5 seconds (9.25s on average). The trial timing is detailed in Fig. 1b. Each trial started with a jittered fixation cross (1-2.5s), followed by the stimulus display. The stimulus was shown at most for 4 seconds. After a hand gesture was recognized, there was another jittered fixation cross (1-2.5s) and a jittered reward feedback (1-2.5s). Following the reward feedback, another fixation cross was displayed for the duration of the difference between 4s and the reaction time plus the processing time for response registration. Eye-tracking data were also collected during both the behavioral and fMRI experiments but is not analyzed herein.

Following the main task, the behavioral and fMRI participants did the same online survey that the M-Turk participants completed to annotate affordance-compatibility scores and familiarity for each object. However, in this phase, the participants annotated only the 48 (or 24 in the fMRI case) stimuli that were used for the main experiment.

### Response decoding

In the behavioral experiment, participants’ hand gestures were tracked by a consumer web camera (30 FPS) and decoded in real-time using a computer vision algorithm, Openpose[82–85], and a fully connected neural network (FCNN) that classifies such movement, frame by frame. For the real-time processing of the video stream, frames were first resized to 192*×*108 pixels and then inputted into BODY 25 model of Openpose with a net resolution setting −1*×*112 and an output resolution −1*×*80. The FCNN classified a participant’s hand position in a frame using an 81-dimensional vector that was created based on Openpose estimations of the 2-dimensional coordinates of 21 key points on the right hand (4 from each finger and 1 from the wrist), as well as the confidence scores of the estimations for each key point.

Specifically, the x and y coordinates of each key point were centered using the wrist location, and then scaled based on the x-directional distance between the left-most and right-most key points, as well as the y-directional distance between the bottom-most and the top-most key points in the current frame. The 81-dimensional input to the FCNN was then built using those centered and scaled x, y coordinates of each key point except for the wrist (20 *×* 2 dimensions), their sum of squares representing the distances between each key point and the wrist (20 dimensions), and confidence scores of the 21 key point estimations (21 dimensions).

The neural network classifier comprised of 3 hidden layers (128, 64, and 32-dimensions, respectively, from low to high layers) and a 4-dimensional softmax output layer in which each unit was associated with one of four hand gestures: pinch, clench, poke, or palm. The deep network classifier was trained using videos of hand motions that were labeled frame-by-frame. The training videos featured hand gestures of 2 males and 2 females, and the total duration of the videos were 40240 frames. We used `MLPClassifier` function from Sklearn 0.23.1 with `lbfgs` optimizer and learning rate 10^*−*5^.

A response in a trial was registered as a valid action only when all frames for 500ms were classified with a probability greater than 95% into one specific hand gesture. To ensure consistency in the starting hand position across trials, it was required that each trial begins with a hand position classified as the palm position by the deep network. The classifier classified responses with 93.2% accuracy and misclassified responses were manually corrected after the experiment (and we then used the correctly decoded action in the subsequent analyses – those *∼*7% errors trials would have produced outcomes with differing probabilities to that intended according to the experimental design, but this variation would not have been noticeable to the participants and was fully accounted for in subsequent data analysis involving the computational models). Reaction times were measured by calculating the time difference between the stimulus onset and the frame where the hand gesture was initiated from the resting palm position using the recorded videos of hand gestures.

In the fMRI experiment, decoding of participant’s actions was found to be much less reliable with the machine-learning algorithm we used successfully in the behavioral study, because participant’s hand position was occluded by the MRI bore. Consequently, we resorted instead to manually decoding each gesture in real-time. Participants’ hand gestures were monitored by the experimenter using a low light USB camera (See3CAM CU30, 40 FPS), and the footage was displayed in the control room in real-time. The experimenter manually but rapidly categorized the movements into one of 3 hand gesture categories (pinch, clench, or poke) thereby enabling participants to interact with the task in real-time. The registered responses were double-checked after each experiment by replaying the recorded video stream, revealing that less than 0.1% of the responses were mistakenly classified during the task performance phase. These error trials were corrected prior to the subsequent data analysis to reflect participant’s intended responses. RTs were calculated using the identical method used for analyzing RTs in the behavioral data.

### Computational modeling of behavior

We tested 9 different models to identify the computational mechanism that best explains the behavioral data. The models were fit to each subject data separately using the Computational Behavioral Modeling (CBM) toolkit `cbm lap` function which calculates model evidence and likelihood of the data using Laplace approximation and estimates parameters using maximum-a-posteriori (MAP) estimation [86]. We used mean 0 and variance 6.25 Gaussian priors for calculating the parameter estimates. Because CBM assumes the parameters to be normally distributed, we applied transformation functions to the parameters: a sigmoid function to model parameters which ranges are between 0 and 1, or an exponential function to model positive parameters. We didn’t use the hierarchical Bayesian inference function of the toolkit because model simulations using the hierarchically estimated parameters couldn’t reproduce the characteristics of the real data. Instead, we performed Bayesian model selection [87] using the log evidence from the CBM outputs to do the Bayesian model comparison. The models were developed solely based on the behavioral data and later tested on the fMRI data.

In order to model affordance perception, we employed the affordance-compatibility score derived from the post-task survey of each participant, which was scaled from 0 to 1. We used individually derived affordance-compatibility scores to fit the model to the behavioral experiment data. However, in the case of fitting the model to each fMRI participant, we used the average scores of annotations from all participants as such a strategy showed a better fit to the data. Notably, the choice of using either individual scores or average scores did not affect the results of the model comparison for either dataset.

### Reinforcement learning model

The reinforcement learning (RL) model is the simplest model we tested that doesn’t consider the effect of affordance, but only models value learning. Action values *Q* for each object are initialized at 0.

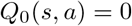

And the probability of choosing an action *a* given a stimulus *s* is a softmax function of action value

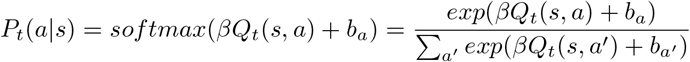

where *β >* 0 is an inverse temperature parameter and *b*_*a*_ *∈* ℝ models action selection bias toward the action independent to the stimulus shown. The action value *Q*_*t*_(*s, a*) for the chosen action *a*_*t*_ to the stimulus *s*_*t*_ at trial *t* is updated based on the delta rule

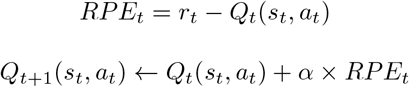

where the reward prediction error (RPE) is the difference between the reward *r*_*t*_ (0 or 1 depending on the realized outcome) at the trial *t* and the action value. 0 *≤ α ≤* 1 is a learning rate parameter. There were 5 free parameters (*β, α, b*_*pinch*_, *b*_*clench*_ and *b*_*poke*_) in this model.

### Bayesian learning model

In this model, instead of using the reinforcement learning rule, value learning is modeled as Bayesian inference [20]. The model tracks the probability of reward of each action given an object using a beta prior and a binomial likelihood. Specifically, the prior on the reward probability of an action *a* to an object *s* is

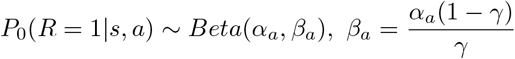

where *α*_*a*_ *>* 0 and 0 *< γ <* 1 are free parameters and the prior mean is *γ*. Given a scenario where an object *s* shown *n* times, with an action *a* being chosen *n*_*a*_ times, resulting in *r*_*a*_ positive outcomes, the posterior reward probability of the action *a* to the object *s* is

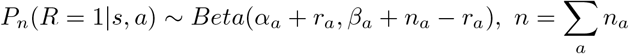

which posterior mean is 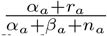.

We then assumed affordance elicits its effect as a bias in the action selection. We used 0-to-1 scaled affordance-compatibility score of action *a* to object *s* from the post-task survey as affordance-compatibility score *Aff* (*s, a*) and used the softmax function,

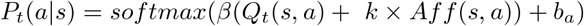

as the action selection policy where *Q*_*t*_(*s, a*) is the posterior mean of *P* (*R* = 1|*s, a*) at trial *t* and *β >* 0, *k >* 0 and *b*_*a*_ *∈* ℝ are free parameters that model inverse temperatures, weight on affordance and action selection bias respectively. In total, this model has 9 free parameters (*α*_*a*_ and *b*_*a*_ for each action type, *γ, β* and *k*)

We also tested models incorporating the affordance into the prior mean instead of including the affordance-based selection bias term in the softmax policy. However, these models exhibited an even poorer fit to the data compared to the model described above.

### Affordance as a prior model

The prior model uses the affordance-compatibility score *Aff* (*s, a*) as the initial action value of an action *a* to an object *s*.

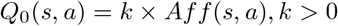

The decision probability and learning rules are identical to the RL model, which results in having one additional free parameter than the RL model (*β, α, b*_*pinch*_, *b*_*clench*_, *b*_*poke*_ and *k*). This model can also be interpreted as implementing a single decision-making system based on affordance and in which the affordance is being relearned based on task experience.

### Affordance as a bias model

The bias model is identical to the RL model except for the calculation of the action selection probability. In the affordance as a bias model, action selection is a softmax function of the linear summation of the action values and the affordance-compatibility scores.

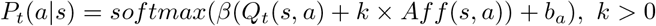

This model has 6 free parameters (*β, α, b*_*pinch*_, *b*_*clench*_, *b*_*poke*_ and *k*).

### Fixed arbitration model

In this model, we assumed there are two different decision-making strategies, value-based decision-making and affordance-based decision-making. Value-based decision-making and value learning is modeled using the RL model previously described.

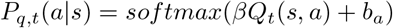

Action selection by the affordance-based decision-making system is modeled as a softmax function of the affordance-compatibility score.

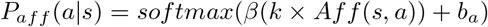

The action selection policy is calculated as a weighted sum of these two policies

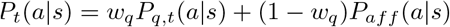

where 0 *≤ w*_*q*_ *≤* 1 is the fixed arbitration weight between the two decision-making strategies. This model has 7 free parameters in total (*β, α, b*_*pinch*_, *b*_*clench*_, *b*_*poke*_, *k* and *w*_*q*_).

### Affordance as a cost model

The cost model is an extension of the fixed arbitration model where we assumed that the response-affordance-incompatibility acts as a cost that is contrasted against the reward signal [88]. Specifically, we modified the delta learning rule to include a cost term which results in a larger reward when an affordance-compatible action is selected so that,

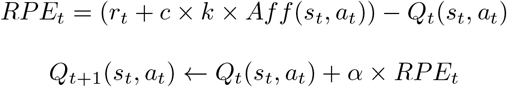

where *c >* 0 is an additional free parameter that models the effect of affordance in learning as a cost. For example, the additional reward will be smaller when the chosen action has a low affordance-compatibility score compared to when it has a high affordance-compatibility which can be interpreted as there was a cost in selecting affordance-incompatible action.

The other components of the model remain identical to the fixed arbitration model, resulting in a cost model with 8 free parameters (*β, α, b*_*pinch*_, *b*_*clench*_, *b*_*poke*_, *k, w*_*q*_ and *c*).

### Conflict-based arbitration model

Like the fixed arbitration model, we assumed that the value-based policy and the affordance-based policy are mixed for the action selection but in this model the arbitration weight dynamically changes trial by trial for each stimulus. The arbitration weight *w*_*q,t*_(*s*) at trial *t* is a logistic function of a conflict variable *c*_*t*_(*s*) which tracks the conflict between the value-based and affordance-based policies given the stimulus *s*. Thus,

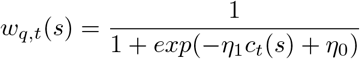

where *η*_1_ *>* 0 and *η*_0_ *∈* ℝ are free parameters. We assume that the conflict signal between the two policies causes an increase in cognitive control which pushes behavior toward value-based decision-making in this paradigm [23, 24]. The conflict variable *c*_*t*_(*s*) is calculated as the square root of the Jensen-Shannon Divergence (JSD) between the value-based and affordance-based policies.

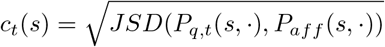

The calculation of value-based and affordance-based polices of the model are identical to the fixed arbitration model which makes this model have 8 free parameters (*β, α, b*_*pinch*_, *b*_*clench*_, *b*_*poke*_, *k, η*_0_ and *η*_1_).

We also tested other design choices such as the energy from a Hopfield network[25] as the proxy of conflict or using a conflict tracking variable that is updated gradually using trial-by-trial JSD. However, the model described above explained the data better than the other tested options.

### Reliability-based arbitration model

This model uses reliabilities [21, 26, 27] of the two strategies as the driving force of arbitration between the value-based and affordance-based decision-making. The reliabilities are calculated using an RL-like updating rule based on unsigned prediction errors [26]. For example, the reliability of value-based decision-making given a stimulus *s*_*t*_ is updated using the following delta rule.

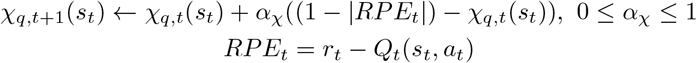

Therefore, the reliability of value-based decision-making will increase when the unsigned RPE is small. Similarly, we defined the reliability of affordance-based decision-making given a stimulus *s*_*t*_ is updated based on

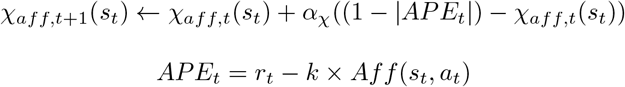

where *APE* is the affordance prediction error and *k* is between 0 to 1 to ensure the APE is between 0 and 1. The reliabilities were initialized to 0. Then the arbitration weight given a stimulus *s* is the logistic function of these two reliabilities. Therefore,

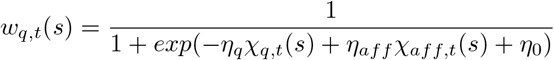

where *η*_*q*_, *η*_*aff*_ *>* 0 and *η*_0_ *∈* ℝ. Identical to the fixed and conflict-based arbitration model, the value-based and affordance-based policies are softmax functions of action values, or affordance-compatibility scores and the final action selection probability is the reliability-based arbitration-weighted sum of the two policies. As a consequence, this model has 10 free parameters (*β, α, b*_*pinch*_, *b*_*clench*_, *b*_*poke*_, *k, α*_*χ*_, *η*_0_, *η*_*q*_ and *η*_*aff*_).

### Performance-based arbitration model

Here, we are proposing a novel arbitration framework that aims to optimize the arbitration weight in terms of maximizing the return. The mathematical derivation of the proposed model is detailed in the supplementary note1. Specifically, we defined value, or performance, of a strategy given a stimulus as the expected returns that can be collected by following that particular strategy in response to the stimulus [18, 22]. Then, the arbitration weight on value-based decision-making given a stimulus *s* is a logistic function of the performances of affordance-based and value-based policies. Thus,

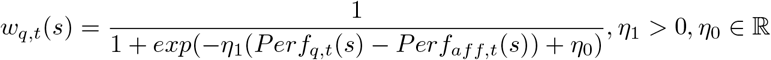

The value-based policy’s performance *Perf*_*q,t*_(*s*) and affordance-based policy’s performance *Perf*_*aff,t*_(*s*) given the stimulus *s* are updated trial by trial by the following delta rules which estimate the performances using inverse propensity scoring from the off-policy policy evaluation literature [28, 29].

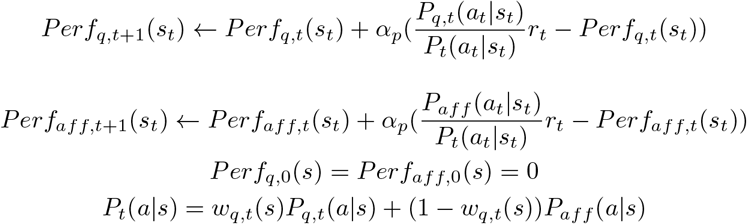

where *r*_*t*_ denotes the realized outcome at the trial *t, P*_*q,t*_(*a*_*t*_|*s*_*t*_) is the probability of selecting the chosen action under the value-based policy at the trial *t*. Similarly, *P*_*aff*_ (*a*_*t*_|*s*_*t*_) represents the probability of selecting the chosen action under the affordance-based policy at the trial t, and *P*_*t*_(*a*_*t*_|*s*_*t*_) denotes the probability of selecting the chosen action. 0 *≤ α*_*p*_ *≤* 1 is a free parameter that denotes the performance updating rate. We called 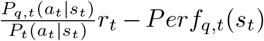, and 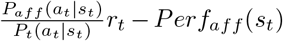 as the performance prediction error of value-based and affordance-based decision-making respectively. Analogous to the other arbitration models, value-based and affordance-based policies are calculated in the same manner as the fixed arbitration model. The final action selection probability is determined by a performance-based arbitration-weighted sum of the two policies. Notably, we set the performance updating rate *α*_*p*_ to be identical to the value learning rate *α*. As a result, the performance-based arbitration model has a total of 8 free parameters (*β, α, b*_*pinch*_, *b*_*clench*_, *b*_*poke*_, *k, η*_0_ and *η*_1_).

### Models with counterfactual value update

We also tested models that update values of unchosen actions. These were implemented using the following learning rule, while keeping the other components the same.

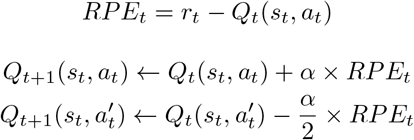

where the reward prediction error (RPE) is the difference between the reward *r*_*t*_ (0 or 1 depending on the realized outcome) at the trial *t* and the action value of chosen action *a*_*t*_ at *t. a*^*′*^ represents the unchosen action at *t* and 0 *≤ α ≤* 1 is a learning rate parameter. The model comparison and simulation results are shown in Supplementary Fig.10

### Model recovery and parameter recovery analyses

We conducted a model recovery analysis across the nine models to ensure that the underlying cognitive mechanism that generated data can be reliably identified using the task design. Using the fMRI version of the task, which has half the number of trials compared to the behavior-only version, each of the nine models was simulated 60 times with the estimated parameters from the real fMRI participants’ data. We then fit the models to each simulated behavior using CBM toolkit and did model comparisons based on log evidence which take into account the number of parameters. Each simulated data was labeled with the best-fitting model, and we calculated the probability of the identified model being the true generative model (Supplementary table 4). It is noteworthy that the probability of the best-fit model being the true generative model for the performance-based and the reliability-based arbitration models were 0.89 and 0.79 each which implies that these models were reliably distinguishable through the model comparison process.

Additionally, we conducted a parameter recovery analysis on the performance-based and the reliability-based arbitration models which were employed for fMRI analyses, to ensure that the model-based variables reliably reflect the underlying cognitive mechanisms. Using the fMRI version of the task which has 480 trials in total, we generated 100 simulated data from each model, with parameters sampled from Gaussian distributions based on sample means and variances of the estimated parameters from the actual fMRI participants data. We estimated the model parameters using CBM toolkit and calculated correlations between the true generative parameters and the estimated parameters. The performance-based arbitration model had a mean Pearson correlation of *r* = 0.76 between true and estimated parameters, while the reliability-based arbitration model had a mean Pearson correlation of *r* = 0.53.

### Behavior general linear model analyses

We conducted mixed-effect GLM analyses on the RT data from the behavior and fMRI studies to examine the effect of choosing affordance-compatible actions on RTs (Figs. 2a,f). As the RT is influenced by both the affordance-compatibility of the response and the value learning process, we incorporated a set of independent variables in the analyses: indicator variables for the affordance-compatibility of the response, whether the response was correct, whether the trial was in a congruent or incongruent condition, whether the trial was in a high or low condition, a categorical variable representing the movement type of the response (i.e., pinch, clench or poke) and a continuous variable representing the trial index. The dependent variable was RT, which was logarithmically transformed to make it normally distributed. The slopes and intercepts were estimated as random effects for each participant and the fixed-effect coefficients are reported in (Supplementary table 1)

In addition, we performed mixed-effect GLM analyses to evaluate the impact of affordance-value congruency on the learning curves (Figs. 2d,e,i,j and figs. 3f,g,l,m). The dependent variable was the difference between congruent and incongruent conditions in terms of frequencies of choosing correct actions, which were extracted for each trial index or the number of rewarded trials so far, for each subject. We included a continuous variable representing the trial index or the number of rewarded trials so far, as regressors in the analyses. The slopes and intercepts were estimated as random effects for each subject and the fixed-effect coefficients are reported in (Supplementary table 3)

We used the `mixedlm` function from the statsmodels 0.12.2 package in Python 3.7.7. The package calculated z statistics of each coefficient by dividing the estimates of coefficients with standard errors. Using the z statistics, the p-values were calculated with respect to a standard normal distribution.

### fMRI data acquisition

fMRI data were acquired at the Caltech Brain Imaging Center (Pasadena, CA), using a Siemens Prisma 3T scanner with a 32-channel radio-frequency coil. The functional scans were conducted using a multi-band echo-planar imaging (EPI) sequence with 72 slices, −30 degrees slice tilt from AC-PC line, 192 mm *×* 192 mm field of view, 2 mm isotropic resolution, repetition time (TR) of 1.12 s, echo time (TE) of 30ms, multi-band acceleration of 4, 54-degree flip angle, in-plane acceleration factor 2, echo spacing of 0.56 ms, and EPI factor of 96. Following each run, both positive and negative polarity EPI-based field maps were collected using similar parameters to the functional sequence, but with a single band, TR of 5.13 s, TE of 41.40 ms, and 90-degree flip angle. T1-weighted and T2-weighted structural images were also acquired for each participant with 0.9 mm isotropic resolution and 230 mm *×* 230 mm field of view. For the T1-weighted scan, TR of 2.55 s, TE of 1.63 ms, inversion time (TI) of 1.15 s, flip angle of 8 degrees, and in-plane acceleration factor 2 were used. The T2-weighted scan was acquired with TR of 3.2 s, TE of 564 ms, and in-plane acceleration factor of 2.

### fMRI data preprocessing

Results included in this manuscript come from preprocessing performed using *fMRIPrep* 20.2.6 [89, 90, RRID:SCR 016216], which is based on *Nipype* 1.7.0 [91, 92, RRID:SCR 002502].

#### Anatomical data preprocessing

A total of 1 T1-weighted (T1w) images were found within the input BIDS dataset.The T1-weighted (T1w) image was corrected for intensity non-uniformity (INU) with `N4BiasFieldCorrection` [93], distributed with ANTs 2.3.3 [94, RRID:SCR 004757]. The T1w-reference was then skull-stripped with a *Nipype* implementation of the `antsBrainExtraction.sh` workflow (from ANTs), using OASIS30ANTs as target template. Brain tissue segmentation of cerebrospinal fluid (CSF), white-matter (WM) and gray-matter (GM) was performed on the brain-extracted T1w using `fast` [95, FSL 5.0.9, RRID:SCR 002823,]. Brain surfaces were reconstructed using `recon-all` [96, FreeSurfer 6.0.1, RRID:SCR 001847,], and the brain mask estimated previously was refined with a custom variation of the method to reconcile ANTs-derived and FreeSurfer-derived segmentations of the cortical graymatter of Mindboggle [97, RRID:SCR 002438,]. Volume-based spatial normalization to one standard space (MNI152NLin2009cAsym) was performed through nonlinear registration with `antsRegistration` (ANTs 2.3.3), using brain-extracted versions of both T1w reference and the T1w template. The following template was selected for spatial normalization: *ICBM 152 Nonlinear Asymmetrical template version 2009c* [98, RRID:SCR 008796; TemplateFlow ID:MNI152NLin2009cAsym].

#### Functional data preprocessing

For each of the 6 BOLD runs per subject (across all tasks and sessions), the following preprocessing was performed. First, a reference volume and its skull-stripped version were generated by aligning and averaging 1 single-band references (SBRefs). A B0-nonuniformity map (or *fieldmap*) was estimated based on two (or more) echo-planar imaging (EPI) references with opposing phase-encoding directions, with `3dQwarp` [99] (AFNI 20160207). Based on the estimated susceptibility distortion, a corrected EPI (echo-planar imaging) reference was calculated for a more accurate co-registration with the anatomical reference. The BOLD reference was then co-registered to the T1w reference using `bbregister` (FreeSurfer) which implements boundary-based registration [100]. Co-registration was configured with six degrees of freedom. Head-motion parameters with respect to the BOLD reference (transformation matrices, and six corresponding rotation and translation parameters) are estimated before any spatiotemporal filtering using `mcflirt` [101, FSL 5.0.9,]. BOLD runs were slice-time corrected to 0.52s (0.5 of slice acquisition range 0s-1.04s) using `3dTshift` from AFNI 20160207 [99, RRID:SCR 005927]. First, a reference volume and its skull-stripped version were generated using a custom methodology of *fMRIPrep*. The BOLD time-series (including slice-timing correction when applied) were resampled onto their original, native space by applying a single, composite transform to correct for head-motion and susceptibility distortions. These resampled BOLD time-series will be referred to as *preprocessed BOLD in original space*, or just *preprocessed BOLD*. The BOLD time-series were resampled into standard space, generating a *preprocessed BOLD run in MNI152NLin2009cAsym space*. First, a reference volume and its skull-stripped version were generated using the custom methodology of *fMRIPrep*. Several confounding time-series were calculated based on the *preprocessed BOLD*: framewise displacement (FD), DVARS and three region-wise global signals. FD was computed using two formulations following Power (absolute sum of relative motions, [102]) and Jenkinson (relative root mean square displacement between affines, [101]). FD and DVARS are calculated for each functional run, both using their implementations in *Nipype* (following the definitions by [102]). The three global signals are extracted within the CSF, the WM, and the whole-brain masks. Additionally, a set of physiological regressors were extracted to allow for component-based noise correction [103, *CompCor*]. Principal components are estimated after high-pass filtering the *preprocessed BOLD* time-series (using a discrete cosine filter with 128s cut-off) for the two *CompCor* variants: temporal (tCompCor) and anatomical (aCompCor). tCompCor components are then calculated from the top 2% variable voxels within the brain mask. For aCompCor, three probabilistic masks (CSF, WM and combined CSF+WM) are generated in anatomical space. The implementation differs from that of Behzadi et al. in that instead of eroding the masks by 2 pixels in BOLD space, the aCompCor masks subtract a mask of pixels that likely contain a volume fraction of GM. This mask is obtained by dilating a GM mask extracted from FreeSurfer’s *aseg* segmentation, and it ensures components are not extracted from voxels containing a minimal fraction of GM. Finally, these masks are resampled into BOLD space and binarized by thresholding at 0.99 (as in the original implementation). Components are also calculated separately within the WM and CSF masks. For each CompCor decomposition, the *k* components with the largest singular values are retained, such that the retained components’ time series are sufficient to explain 50 percent of variance across the nuisance mask (CSF, WM, combined, or temporal). The remaining components are dropped from consideration. Head-motion estimates calculated in the correction step were also placed within the corresponding confounds file. The confound time series derived from head motion estimates and global signals were expanded with the inclusion of temporal derivatives and quadratic terms for each [104]. Frames that exceeded a threshold of 0.5 mm FD or 1.5 standardised DVARS were annotated as motion outliers. All resamplings were performed with *a single interpolation step* by combining all the pertinent transformations (i.e. head-motion transform matrices, susceptibility distortion correction when available, and co-registrations to anatomical and output spaces). Gridded (volumetric) resamplings were performed using `antsApplyTransforms` (ANTs), configured with Lanczos interpolation to minimize the smoothing effects of other kernels [105]. Non-gridded (surface) resamplings were performed using `mri vol2surf` (FreeSurfer).

Many internal operations of *fMRIPrep* use *Nilearn* 0.6.2 [106, RRID:SCR 001362], mostly within the functional processing workflow. For more details of the pipeline, see the section corresponding to workflows in *fMRIPrep*’s documentation.

### Whole brain general linear model analysis

FSL (FMRIB Software Library) 6.0 software package was used to analyze the fMRI data. Following the preprocessing using fMRIPrep, the fMRI data was convolved with a 3D Gaussian smoothing kernel (8.0mm FWHM) using FSL SUSAN. Subsequently, BOLD signals for each voxel were scaled to ensure that the mean BOLD signal of each voxel is 100. Before running GLM analyses, first 4 volumes were discarded and a high-pass filter with a cutoff 128s was applied. Framewise displacement, global signal, global signal derivative, 6 head-motion parameters (x, y, z translations and rotations), and temporal CompCor from fMRIPrep were included as nuisance variables in every analysis. Stick regressors that represent onsets of stimulus, fixation cross and outcome were also included in the GLMs. The response, or the hand movement, was modeled using a parametric regressor detailed in the following section. We used the fixed-effect higher-level modeling for individual-level inference that averages across blocks and FSL FLAME1 for group-level inference.

### Hand movement parametric regressor

In order to minimize potential confounding effects that may arise from using natural hand gestures as the action in the experiment, we designed a hand movement regressor based on 2-dimensional 21 key points for the right hand extracted from the hand video using Openpose. This regressor was included in all of the analyses.

First, video frames were resized into 162*×*108 pixels to fit the Openpose net resolution −1*×*122. By utilizing the BODY 25 model from the Openpose, key points were extracted frame by frame with the output resolution setting −1*×*80. Next, the key points were weighted-moving-averaged with a window of 20 frames (about 500ms) to reduce the noise stemming from Openpose’s estimation error. The confidence scores of Openpose estimations were utilized as the weights in order to mitigate the impact of uncertain estimations. Then, the first principal component of the weighted-moving-averaged 42-dimensional data (21*×*2) was calculated and used as the parametric regressor that models the hand movement. In all of our GLM analyses, the hand movement regressor consistently identified the left primary motor cortex, left premotor cortex, left primary somatosensory cortex, and bilateral inferior parietal lobule as the regions associated with the right-hand movement (Supplementary Fig. 14).

### Identifying decision-making regions

A GLM analysis was conducted to investigate the regions associated with the affordance-based and value-based decision-making, as well as the action selection which is the integration of those two systems (GLM1). The chosen action’s affordance-compatibility score (*Aff* (*s*_*t*_, *a*_*t*_)), chosen action’s value (*Q*_*t*_(*s*_*t*_, *a*_*t*_)), the action selection probability of chosen action (*P*_*t*_(*a*_*t*_|*s*_*t*_)) from the performance-based arbitration model models were included as parametric regressors at the onsets of stimuli. A parametric regressor of the reward prediction error was also included at the onsets of outcomes. Another GLM analysis using model-based parameters from the reliability-based arbitration model was conducted as well (GLM2). The results were cluster corrected with a cluster-defining threshold of *z* = 3.1.

### Identifying arbitration regions

Two GLM analyses were conducted to identify regions related to arbitration between value-based and affordance-based decision-making. One GLM analysis utilized arbitration-related parameters obtained from the performance-based arbitration model (GLM3) and the other one used those from the reliability-based arbitration model (GLM4).

GLM3 included performances of the value-based and affordance-based decision-making systems (*Perf*_*q,t*_(*s*_*t*_), *Perf*_*aff,t*_(*s*_*t*_)), and the difference between those two (*Perf*_*aff,t*_(*s*_*t*_)*−Perf*_*q,t*_(*s*_*t*_)) as parametric regressors at the onsets of stimuli. Parametric regressors of the performance prediction errors of value-based and affordance-based policies each were also included at the onsets of outcomes in the GLM3.

In GLM4, reliabilities of the value-based and affordance-based decision-making systems (*χ*_*q,t*_(*s*_*t*_), *χ*_*aff,t*_(*s*_*t*_)), and the difference between those two (*χ*_*aff,t*_(*s*_*t*_) *− χ*_*q,t*_(*s*_*t*_)) were included as parametric regressors at the onsets of stimuli. A parametric regressor of the affordance prediction error was included as well at the onsets of outcomes in the GLM4. The results were cluster corrected with a cluster-defining threshold *z* = 3.1.

### Neural correlates of affordance-incompatible action execution

To identify neural correlates that are activated more when executing affordance-incompatible actions compared to affordance-compatible actions [11, 12], a model-free GLM analysis was conducted (GLM5). GLM5 included stick regressors representing the stimuli identities at the stimulus onsets, 2 regressors representing whether the affordance and response were compatible or not at the response onsets, and a parametric regressor of reward at the outcome onsets. While the contrast between the affordance response compatibility regressors have not survived the cluster correction with a cluster-defining threshold *z* = 3.1, Supplementary Fig. 14g shows the uncorrected results with a threshold set at *p <* 0.001, corresponding to *z ≥* 3.1.

### Neural representation strength analysis

To calculate the extent to which the identified regions from GLM analyses can be explained by the corresponding cognitive variables, we computed the mean z-statistics for those specific variables across the voxels within the corresponding group-level regions. The regions were defined using the survived clusters from GLM1,2,3 and 4 (For the details, see Fig. 4 and Supplementary Fig. 12). We then compared the mean z-statistics for each individual with the frequency of choosing the most rewarding action.

Similarly, the increased BOLD activity for executing affordance-incompatible actions compared to executing affordance-compatible actions was calculated using the contrast between the affordance-response compatibility regressors from GLM5. The z-statistics of the contrast were averaged across the voxels within functionally-defined regions of interest (ROI). The ROIs were defined in the same way as in the previous analyses. We then compared the average z-statistics for each individual with the frequency of choosing the affordance compatilble action or the reaction time effect which is the relative increase in reaction time when choosing affordance-incompatible actions compared to affordance-compatible actions 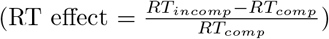.

## 5 Acknowledgments

We thank Shinsuke Shimojo, Jeff Cockburn, Vincent Man and Seungyong Moon for discussion and suggestions, Weilun Ding, Thomas Henning, Seokyoung Min, Jihong Min, Areum Kim, HyeongChan Jo, and Serim Ryou for their help in implementing the task. **Funding:** This work was supported by a graduate innovator grant award from Caltech’s Tianqiao and Chrissy Chen Institute for Neuroscience to S.Y. **Author contributions:** S.Y. and J.P.O. conceived and designed the study, S.Y. performed experiments and S.Y. and J.P.O. analyzed and discussed results. S.Y. and J.P.O. wrote the manuscript. **Competing interests:** The authors declare that they have no competing interests. **Data and materials availability:** The data, code and analysis results utilized in this manuscript will be available online at the time of publication.

## 6 Supplementary information

Supplementary Note1 and 2

Figs. S1 to S16

Tables S1 to S7

