## Supplementary information for "Computational and neural mechanisms underlying the influence of action affordances on value learning"

### 1 Supplementary information

**Note1: Derivation of performance-based arbitration model**
The objective of the performance-based arbitration model in the context of the given research paradigm, or for general contextual bandit problems, is to maximize the return with respect to the arbitration weight. More precisely, we consider a scenario in which a behavior policy, denoted as $\pi$, is a linear mixture of multiple component policies, represented as $\pi_i$.

$$\pi(a|s) = \sum_i w_i(s)\pi_i(a|s), \ \sum_i w_i = 1, \ 0 \leq w_i \leq 1 \tag{1}$$

In our experiment, we assumed the final action selection probability corresponds to the behavior policy and the value-based and affordance-based decision-makings are the component policies.

If we define a value, or performance, of a component policy $\pi_i$ with respect to a given context, or stimulus, $s$ as the expected return obtained by following the policy $\pi_i$ when the stimulus $s$ is shown:

$$Perf_{\pi_i}(s) := \sum_a \pi_i(a|s)R(s,a) \tag{2}$$

Then the performance of the policy $\pi$ at the state $s$ can be decomposed into the linear summation of the performances of the component policies.

$$Perf_{\pi}(s) := \sum_a \pi(a|s)R(s,a) = \sum_a \sum_i w_i(s)\pi_i(a|s)R(s,a) \tag{3}$$

$$= \sum_i w_i(s) \sum_a \pi_i(a|s)R(s,a) = \sum_i w_i(s)Perf_{\pi_i}(s) \tag{4}$$

Therefore, determining arbitration weights in terms of maximize the performance of the behavior policy $\pi$ can be formulated as identifying one of the component policies that yields the highest performance and assigning weight of 1 to it.

$$max_{w_i(s)}(Perf_{\pi}(s)) = max_i(Perf_{\pi_i}(s)) \tag{5}$$

Therefore, the arbitration problem can be solved if the performances of the component policies can be estimated. Given a trajectory of actions, and outcomes generated by the behavior policy $\pi$ at a state $s$, the approximation of the performance of a component policy $\pi_i$ can be achieved by employing various methods from the off-policy evaluation literature. One such method is the inverse propensity scoring estimator [1, 2], which can be written in the following form:

$$Perf_{\pi_i}(s) \approx \frac{1}{T} \sum_{t=1}^{T} \frac{\pi_i(a_t|s)}{\pi(a_t|s)} r_t \tag{6}$$

where $T$ is the number of trials at the state $s$ in the trajectory and $r_t$ is the realized outcome at the $t$-th trial at the state $s$. In our scenario of having a value-based policy as one of the component policies, the behavior policy and the value-based policy are nonstationary and get updated every trial. Therefore, it is more practical to assign greater weight to recent events while estimating the performance of component policies. In this context, the above equation can be approximated trial by trial by the following delta rule.

$$Perf_{\pi_i,t+1}(s_t) \leftarrow Perf_{\pi_i,t}(s_t) + \alpha_p(\frac{\pi_i(a_t|s_t)}{\pi(a_t|s_t)}r_t - Perf_{\pi_i,t}(s_t)),\ 0 \leq \alpha_p \leq 1 \tag{7}$$

where we called $(\frac{\pi_i(a_t|s_t)}{\pi(a_t|s_t)}r_t - Perf_{\pi_i,t}(s_t))$ the performance prediction error of $\pi_i$ and $s_t$ is the stimulus shown at the trial $t$. However, in practical scenarios, accurately determining the performance of each component policy is often unfeasible. Instead, the optimal weighting can be approximated by weighting each component policy with the probability that the policy's performance is larger than any other component policies.

$$max_{w_i(s)}(Perf_\pi(s)) \approx \sum_i P(Perf_{\pi_i}(s) > Perf_{\pi_j}(s), \forall j \neq i)Perf_{\pi_i}(s) \tag{8}$$

The ideal approach for calculating the probability $P(Perf_{\pi_i}(s) > Perf_{\pi_j}(s), \forall j \neq i)$ would involve approximating the full distribution of the performance of each component policy which can be implemented using, for example, distributional RL [3]. However, a simpler approach to approximate such probability would be to employ a softmax function of the performances estimated by the learning rule (Eq. 7) which approximates means of performances.

It's noteworthy that the above framework for the multi-armed bandit problem can be extended to a general Markov decision process. For example, the performance of a component policy $\pi_i$ at a state s can be defined as the expected return by executing $\pi_i$ at the state s and following $\pi$ thereafter.

$$Perf_{\pi_i}(s) := \sum_a \pi_i(a|s)(R(s,a) + \gamma \sum_{s'} P(s'|s,a)Perf_\pi(s')) \tag{9}$$

$$Perf_\pi(s) := \sum_a \pi(a|s)(R(s,a) + \gamma \sum_{s'} P(s'|s,a)Perf_\pi(s')) = V_\pi(s) \tag{10}$$

where $P(s'|s,a)$ is a state transition probability from the current state s to the subsequent state $s'$ after selecting an action $a$, and $\gamma$ is a discounting rate. Then, the performance of the behavior policy can be expressed as the linear summation of component policies' performances.

$$Perf_\pi(s) = \sum_i w_i(s)Perf_{\pi_i}(s) \tag{11}$$

As a result, the problem of maximizing the behavior policy's performance with respect to arbitration weights can be reduced to the task of identifying the component policy with the highest performance.

Because $\sum_j w_j(s')Perf_{\pi_j}(s') \approx max_j Perf_{\pi_j}(s')$, the performance of $\pi_i$ can be written in Bellman equation form:

$$Perf_{\pi_i}(s) = \sum_a \pi_i(a|s)(R(s,a) + \gamma \sum_{s'} P(s'|s,a) \sum_j w_j(s')Perf_{\pi_j}(s')) \quad (12)$$

$$\approx \sum_a \pi_i(a|s)(R(s,a) + \gamma \sum_{s'} P(s'|s,a) max_j Perf_{\pi_j}(s')) \quad (13)$$

Analogous to Eq. 7, the performances of component policies can be estimated by using the following delta rule:

$$Perf_{\pi_i,t+1}(s_t) \leftarrow Perf_{\pi_i,t}(s_t) + \alpha_p(\frac{\pi_i(a_t|s_t)}{\pi(a_t|s_t)}(r_t + \gamma max_j Perf_{\pi_j,t}(s_{t+1})) - Perf_{\pi_i,t}(s_t)) \quad (14)$$

**Note2: Additional model simulation results**

In the main text, we demonstrated that the learning slope in the low conditions is not steeper than it in the high conditions when the learning slopes were plotted as the function of the number of rewarded trials previously encountered. To explore this observation further, a fixed-arbitration model incorporating two distinct learning rate parameters was fit to the data. This additional analysis also revealed no significant difference in learning rates between high and low conditions ($t(18) = -1.57, p = 0.13$ in the behavioral, $t(29) = -0.47, p = 0.65$ in the fMRI data).

There is a potential that participants adopted a simpler strategy, such as win-stay lose-shift, instead of value learning. However, this possibility can be discounted based on the estimated learning rates derived from model fitting. If participants were using a win-stay lose-shift strategy, the estimated learning rates would be expected to be near 1. In contrast, the learning rates obtained using the performance-based model were $0.227 \pm 0.206$ and $0.269 \pm 0.173$ for the behavioral and fMRI data, respectively.

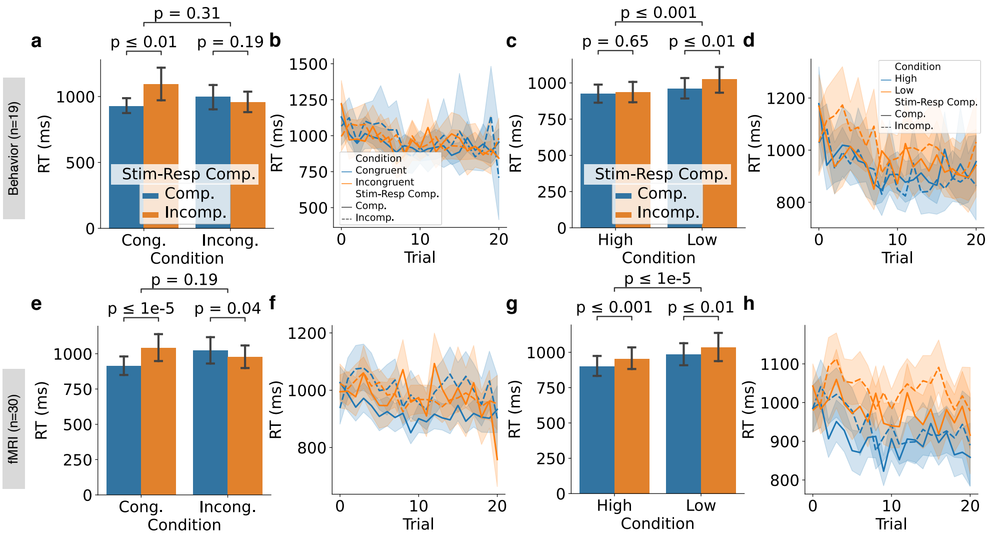


**Supplementary Fig. 1**: Effects of stimulus-response compatibility and value learning on reaction times. (a and e) The RTs for making affordance-compatible actions were significantly faster in the congruent condition, whereas the trend was in the opposite direction in the incongruent condition. This is because affordance-compatible actions have higher action values in the congruent condition, but not in the incongruent condition (paired $t$ tests, $t(18) = -3.30$ and $t(18) = 1.36$ for comparisons within congruent and incongruent conditions respectively in the behavioral, $t(29) = -5.87$ and $t(29) = 2.17$ for the same comparisons in the fMRI experiment). But there was no significant overall RT difference on average between congruent and incongruent conditions (paired $t$ tests, $t(18) = 1.02$ and $t(29) = -1.32$ for the behavioral and fMRI experiments respectively). (b and f) Similar to plots A and E, but the RTs were averaged across blocks. Across learning, as action value learning progresses, RTs become faster. (c and g) RT contrasts plotted similarly to plots A and E but now examined separately within high and low value conditions. The RTs were slower in the low value condition compared to the high value condition (paired $t$ tests, $t(18) = -3.79$ and $t(29) = -7.60$ for the behavioral and fMRI experiments respectively). Also, RTs for executing affordance compatible actions were faster in general regardless of the value manipulation (paired $t$ tests, $t(18) = -0.46$ and $t(18) = -3.58$ for comparisons within high and low conditions respectively in the behavioral, $t(29) = -3.99$ and $t(29) = -3.07$ for the same comparisons in the fMRI experiment). (d and h) Analogous to B and F, the RTs depicted in plots C and G were averaged across blocks.

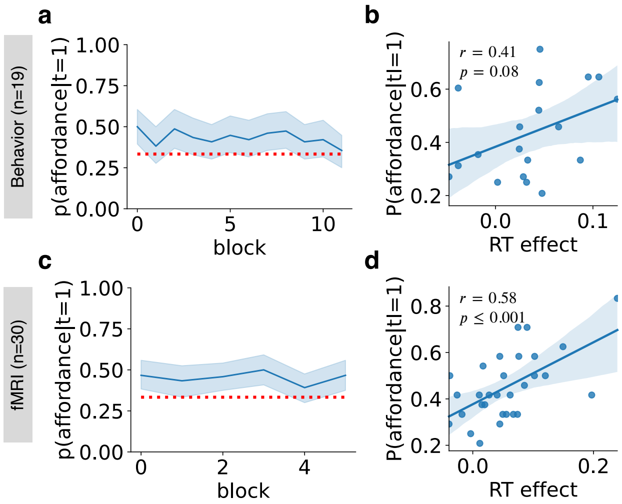


**Supplementary Fig. 2**: Behavioral effects of affordance during value learning and decision making. Upper-row plots depict results from the behavioral study while the lower-row plots are from the fMRI study. (a and c) Initial action selection bias toward the affordance-compatible action over blocks. The red dotted lines show the chance level of 1/3. The mixed linear regression of initial selection bias on block index showed non-significant slopes in both datasets (Fixed-effect coefficients for the slope: $\beta$ = -0.005, z=-1.115 and p=0.265 in the behavioral data, $\beta$=-0.002, z=-0.232 and p=0.817 in the fMRI data).(b and d) The additional RT for executing an affordance-incompatible action relative to an affordance-compatible action, or RT effect, is positively correlated with the participant's tendency to select the affordance-compatible action as the initial response to each object.

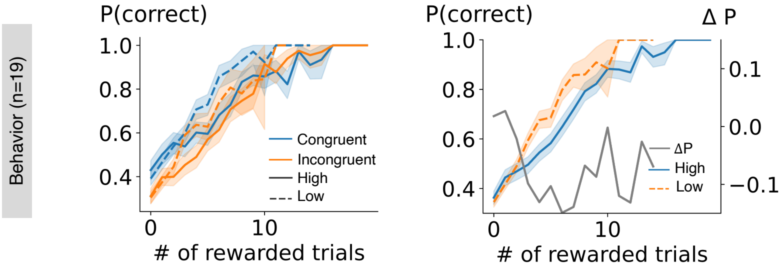


| | Coef. | Std. Err. | z | P>lzl |
|---|---|---|---|---|
| **Intercept** | -0.006 | 0.026 | -0.218 | 0.828 |
| **Number of rewarded trials** | -0.006 | 0.005 | -1.295 | 0.195 |

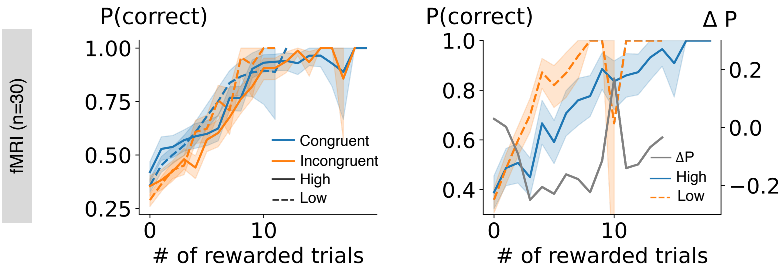


| | Coef. | Std. Err. | z | P>lzl |
|---|---|---|---|---|
| **Intercept** | -0.024 | 0.077 | -0.316 | 0.752 |
| **Number of rewarded trials** | -0.010 | 0.016 | -0.656 | 0.512 |

**Supplementary Fig. 3**: Learning slopes by conditions and their differences between high and low conditions. The table shows the fixed-effect coefficients obtained from the mixed-effect GLM analysis on the difference in the choice accuracy between high and low conditions, with the number of rewarded trials previously encountered being the regressor. $\Delta P = P(correct|high) - P(correct|low)$

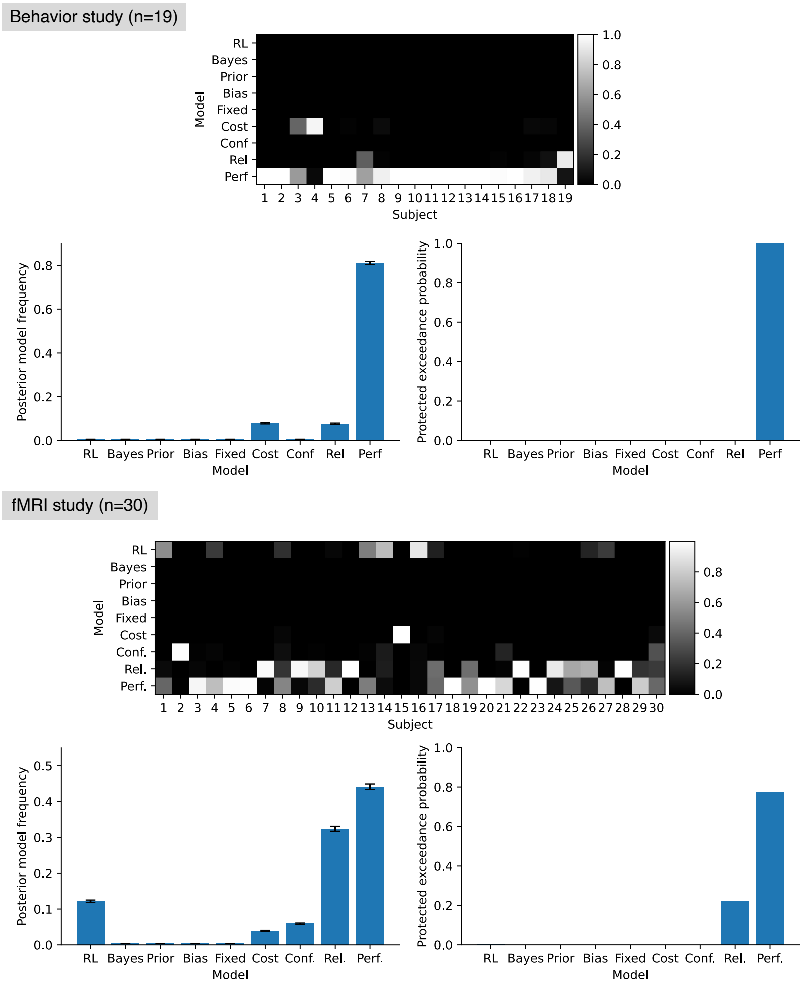


**Supplementary Fig. 4**: Bayesian model selection results. The upper panel displays the posterior probability of each model to best explain each participant. The lower-left panel shows model frequencies over models, while the lower-right panel shows protected exceedance probabilities of models. Overall, the performance-based arbitration model best fit the datasets.

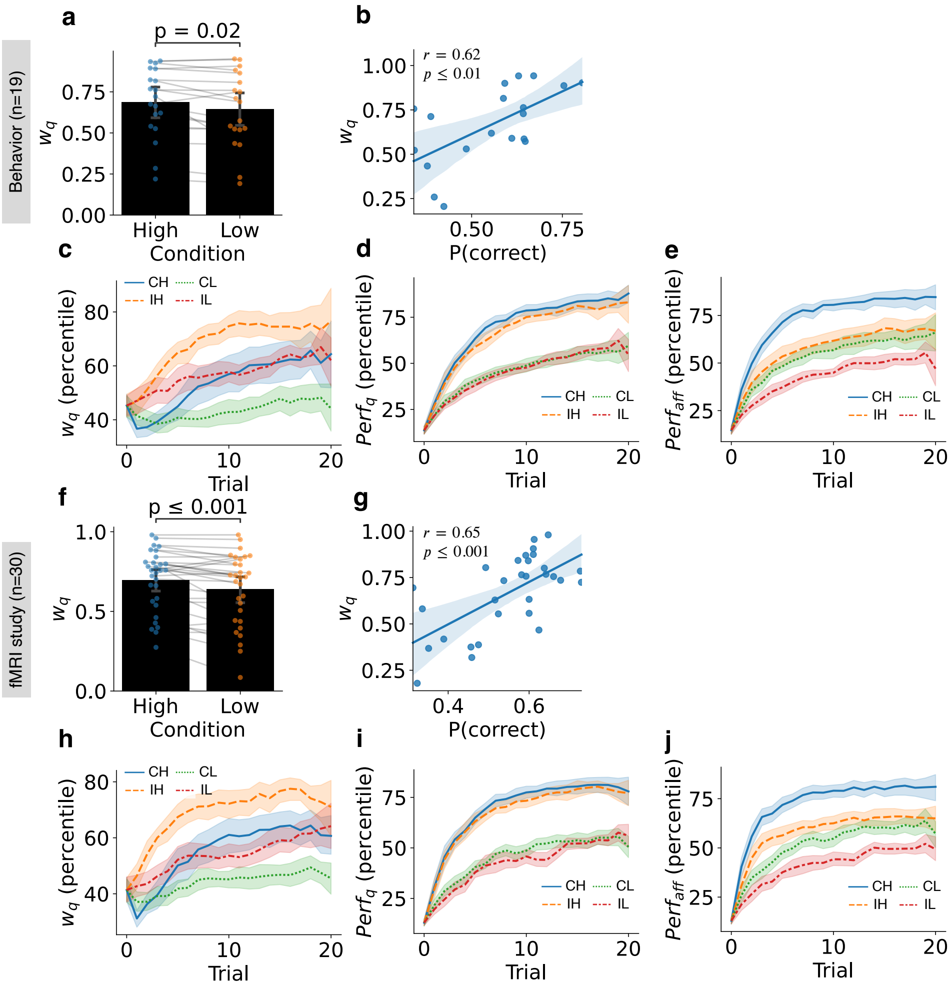


**Supplementary Fig. 5**: Arbitration variables extracted from the performance-based arbitration model. (a and f) Modulation of arbitration by the reward probability manipulation. Arbitration weights are calculated using the performance-based arbitration model for each trial and averaged for each participant and condition. The results suggest that value-based decision-making is more favored in high-value conditions (paired $t$ tests, $t(18) = 2.53, t(29) = 3.77$ each). (b and g) Participants who have a larger average weight toward the value-based decision-making across trials exhibited better accuracy in choosing the most rewarding action. (c and h) Modulation of the performance-based arbitration weight over trials by conditions. CH: congruent high; CL: congruent low; IH: incongruent high; IL: incongruent low. The arbitration weights were transformed into the percentiles within each subject. (d and i) Modulation of the performance of the value-based decision-making system over trials by conditions. (e and j) Modulation of the performance of the affordance-based decision-making system over trials by conditions. The performances were transformed into the percentiles within each subject. All the curves were averaged across blocks and the error-bar shows 95% interval of estimated statistics.

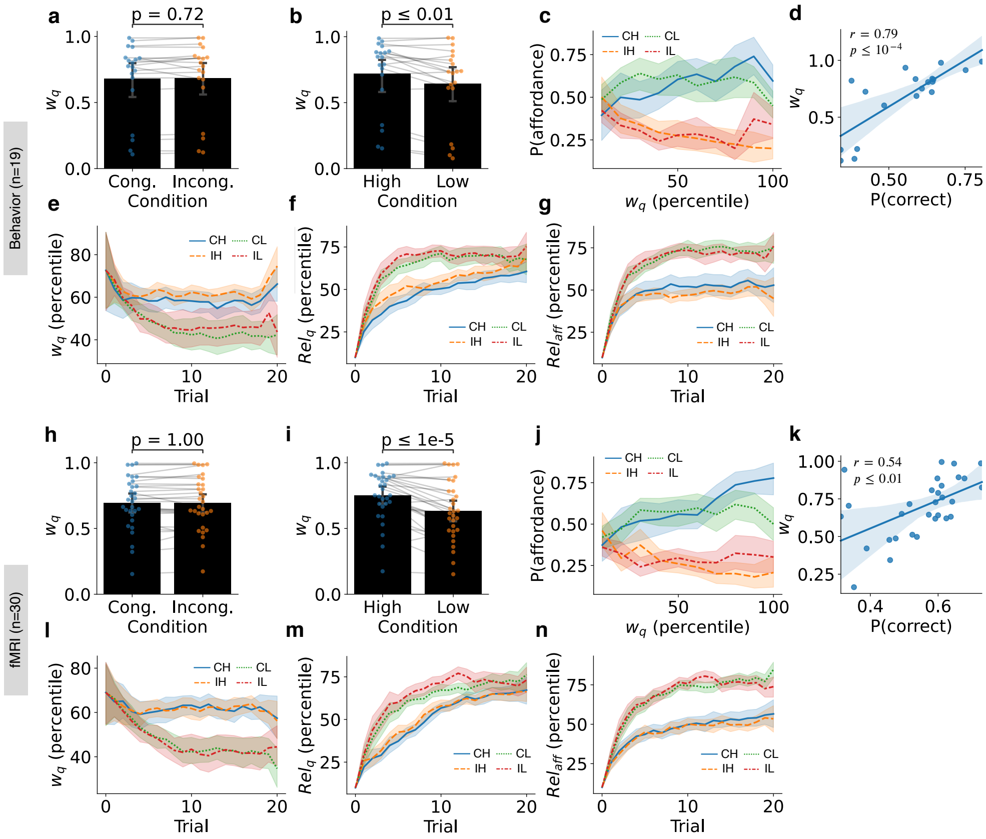
Behavior (n=19)
fMRI (n=30)
a
p = 0.72
$w_q$
Cong.
Incong.
Condition
b
p ≤ 0.01
High
Low
Condition
c
P(affordance)
$w_q$ (percentile)
CH
CL
IH
IL
d
$r = 0.79$
$p \leq 10^{-4}$
P(correct)
e
$w_q$ (percentile)
Trial
f
$Rel_q$ (percentile)
g
$Rel_{aff}$ (percentile)
h
p = 1.00
i
p ≤ 1e-5
j
k
$r = 0.54$
$p \leq 0.01$
l
m
n

**Supplementary Fig. 6**: Arbitration variables extracted from the reliability-based arbitration model. (a and h) Modulation of arbitration by affordance-value congruency. Arbitration weights were calculated using the reliability-based arbitration model for each trial and averaged for each participant and condition. The congruency between affordance and value didn't affect the reliability-base arbitration weight. (paired $t$ tests, $t(18) = 0.36, t(29) = 0.00$ each) (b and i) Modulation of arbitration by the reward probability manipulation. These results suggest that value-based decision-making is more favored in the high-value condition. (paired $t$ tests, $t(18) = 3.86, t(29) = 5.44$ each) (c and j) Frequency of choosing affordance-compatible actions in the data as a function of the arbitration weight on value-based decision-making extracted from the reliability-based arbitration model. The arbitration weights were transformed into the percentiles within each participant. The results show that in the trials when the model preferred affordance-based decision-making, the actual choices were more biased toward affordance-compatible actions. Notably, this trend was only evident in incongruent conditions as the responses based on affordance and value were indistinguishable in congruent conditions. CH: congruent high; CL: congruent low; IH: incongruent high; IL: incongruent low. (d and k) Participants who have a larger average weight toward value-based decision-making across trials exhibited better accuracy in choosing the most rewarding action. (e and l) Modulation of the reliability-based arbitration weight over trials by conditions. The arbitration weights were transformed into percentiles within each subject. (f and m) Modulation of reliability of the value-based decision-making system over trials by conditions. (g and n) Modulation of reliability of the affordance-based decision-making system over trials by conditions. The reliabilities were transformed into the percentiles within each subject. All the curves were averaged across blocks and the error-bar shows 95% interval of estimated statistics.

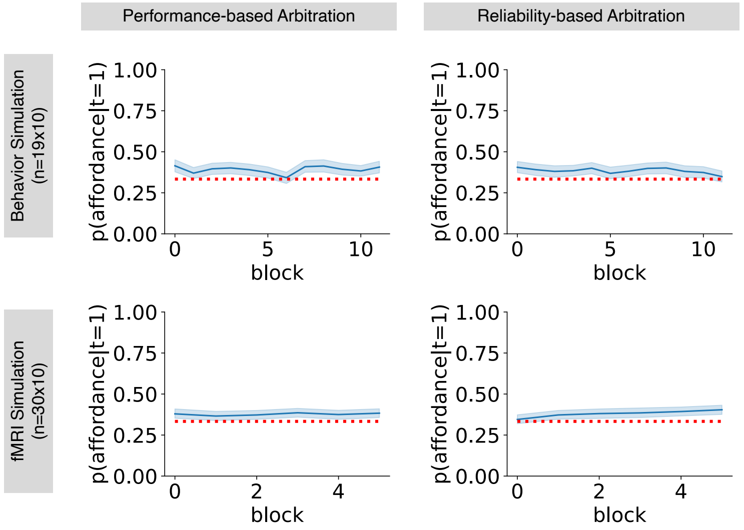


**Supplementary Fig. 7**: Simulated initial action selection bias toward the affordance-compatible action over blocks. The red dotted lines show the chance level 1/3.

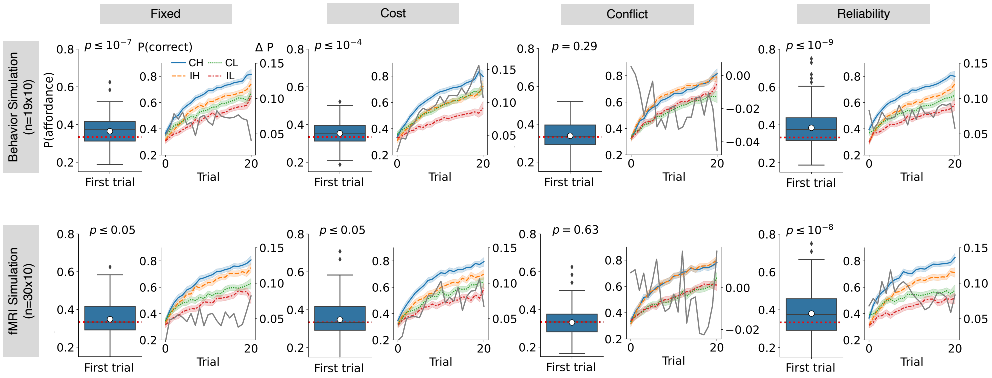


**Supplementary Fig. 8**: Simulated behaviors generated by various computational models. Each plot corresponds to Fig. 3e,k or 3f,l in which behaviors were generated by the computational model labeled on the top of each column. The t statistics of the initial response bias toward affordance compatible actions are $t(189) = 5.56, 4.07, 1.06,$ and $6.59$ each for the simulations based on the behavioral experiment, $t(299) = 2.55, 2.45, -0.48,$ and $6.00$ each for the simulations based on the fMRI experiment.

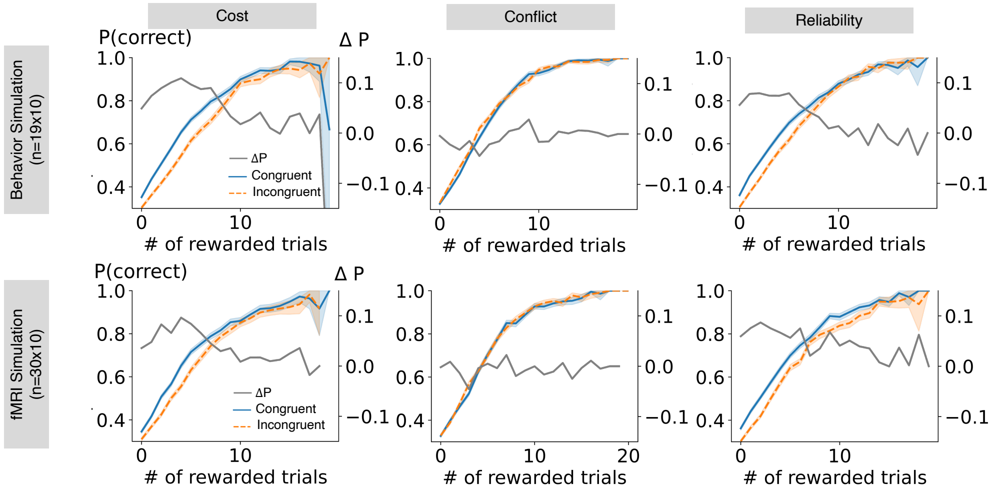


**Supplementary Fig. 9**: Simulated learning slopes and their differences produced by various computational models. Each plot corresponds to Fig. 3g or 3m in which behaviors were generated by the computational model labeled on top of each column.

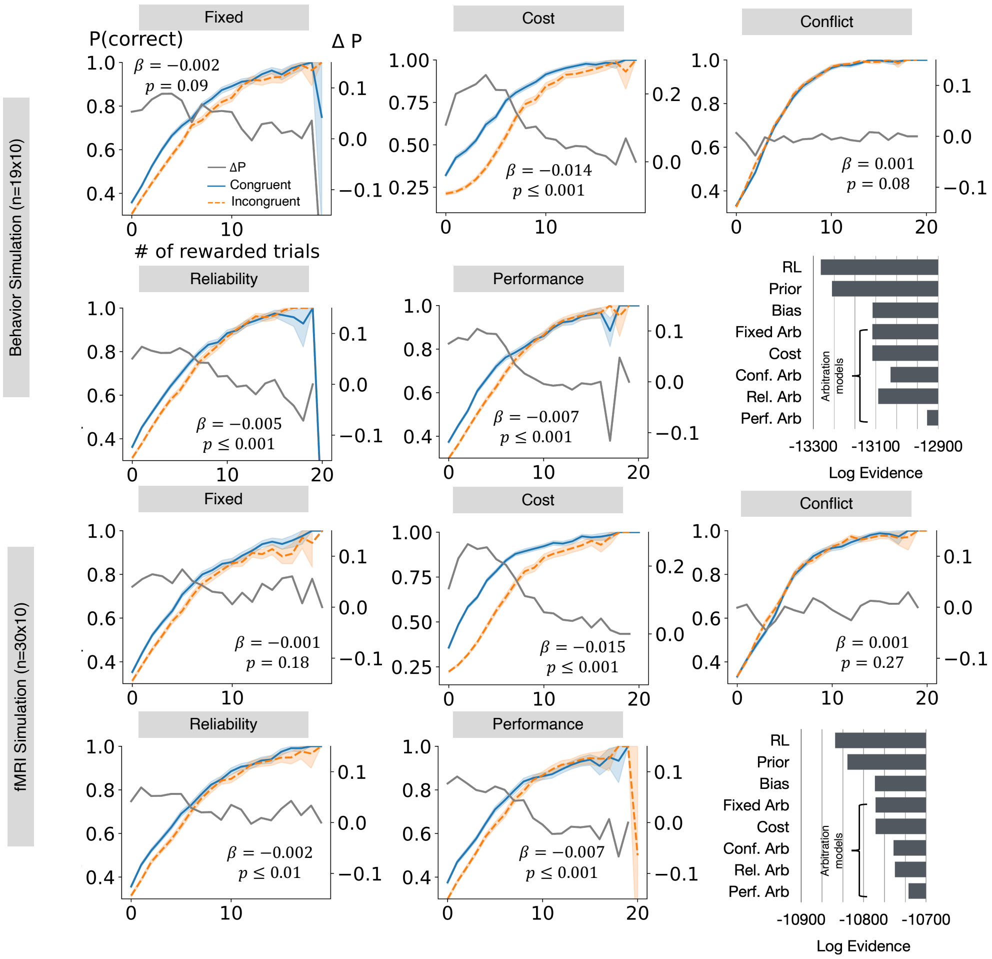


**Supplementary Fig. 10**: Simulated learning slopes and their differences produced by computational models that includes value updates for unchosen actions. Each plot corresponds to Fig. 3g or m in which behaviors were generated by the computational model labeled on top of each column. The $\beta$ and $p-$value represent the fixed-effect coefficient of the slope of the difference between learning slopes and its corresponding $p-$value. The bar graphs represent the model fitting results for models incorporating value updates in unchosen actions. The simulation results from models with unchosen action value updates were generally comparable with the model simulation results without the unchosen action value update, except for the cost model which exhibited much steeper learning slope differences and different ranges for both the learning slopes and their differences compared to the actual data shown in Fig. 2e or j

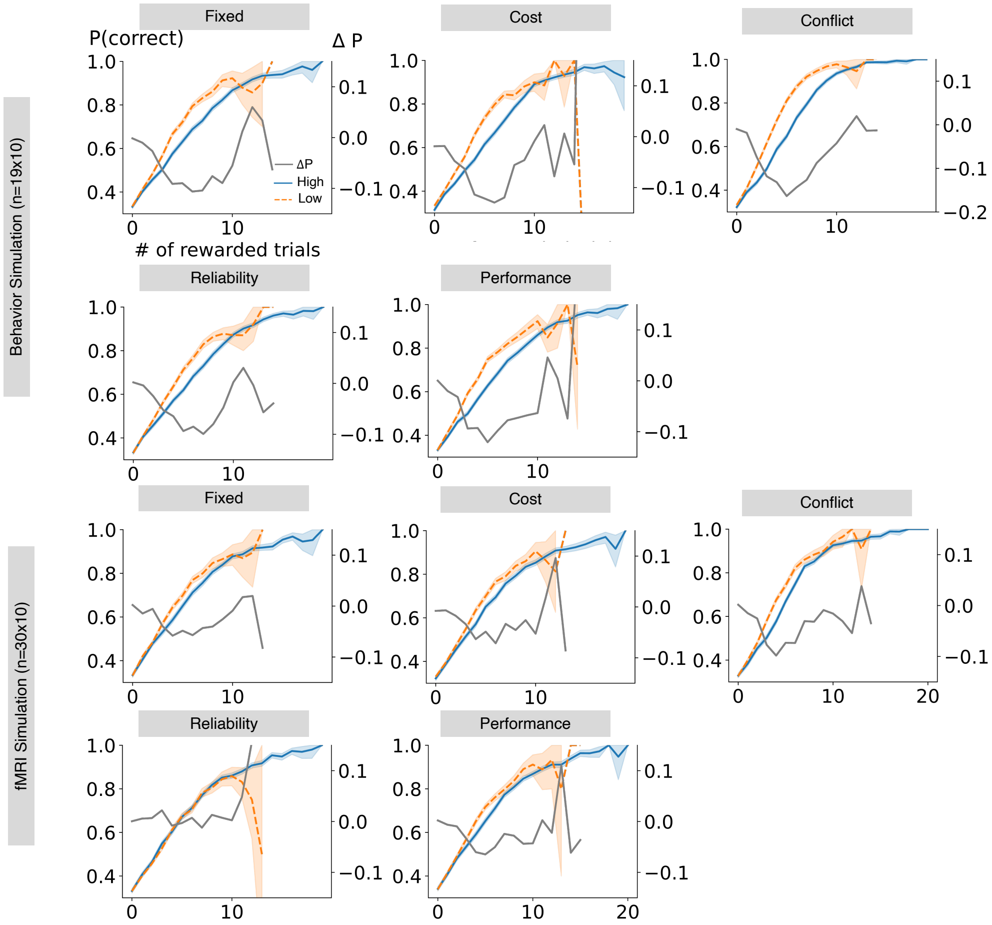


**Supplementary Fig. 11**: Simulated learning slopes and their differences produced by various computational models. Each plot corresponds to Supplementary Fig.3 in which behaviors were generated by the computational model labeled on top of each subplot.

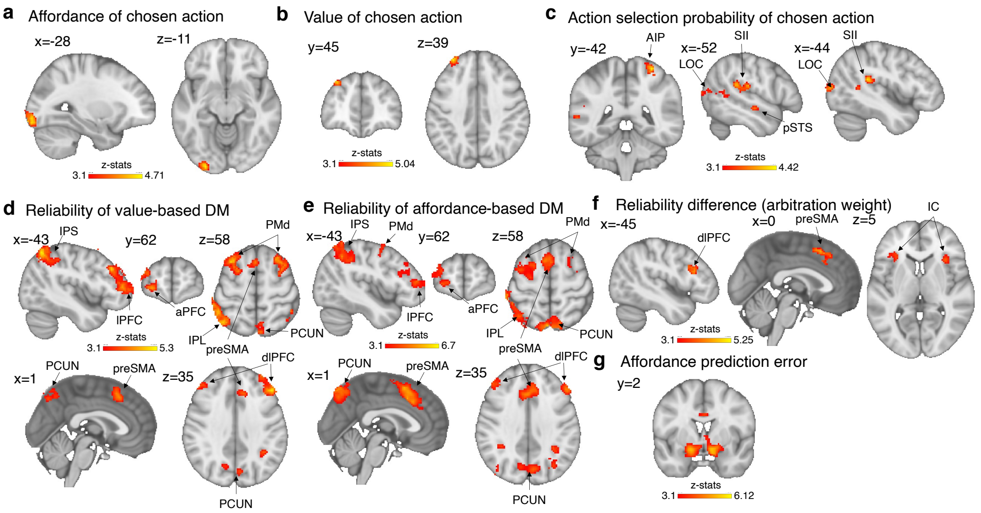


**Supplementary Fig. 12**: Neural implementation of the reliability-based arbitration model. (a) Affordance-compatibility scores of the chosen action correlated with the high-level ventral visual stream including V3 and V4 in the left occipital lobe (b) Action values from the reliability-based arbitration model correlated with activity in the dlPFC (c) Action selection probabilities of the chosen action from reliability models correlated with activity in AIP, SII, pSTS and LOC. (d and e) Reliabilities of both decision-making systems significantly correlated with the preSMA, IPS, lPFC, aPFC, PMd, PCUN, and IPL. (f) The difference between the reliabilities of the two systems ($Rel_{aff} - Rel_q$), which is directly related to the arbitration weight, was identified in the preSMA, dlPFC, and IC. (g) Affordance prediction error (APE) signals which are putatively employed for tracking the reliability of the affordance-based decision-making system were identified in the ventral striatum. All the results were cluster-corrected $p < 0.05$ with the cluster defining threshold $z = 3.1$.

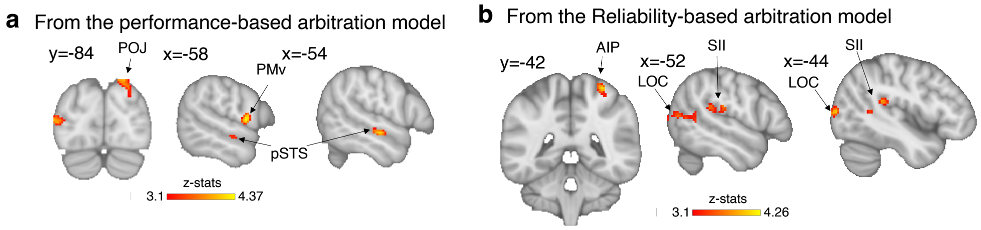


**Supplementary Fig. 13**: Neural correlates of action selection probability of chosen action after controlling reaction times. (a) Results from GLM1 with RT (uncorrected with the threshold $p < 0.001$). (b) Reuslts from GLM2 with RT (cluster corrected with the threshold $p < 0.05$ with the cluster defining threshold $z = 3.1$.)

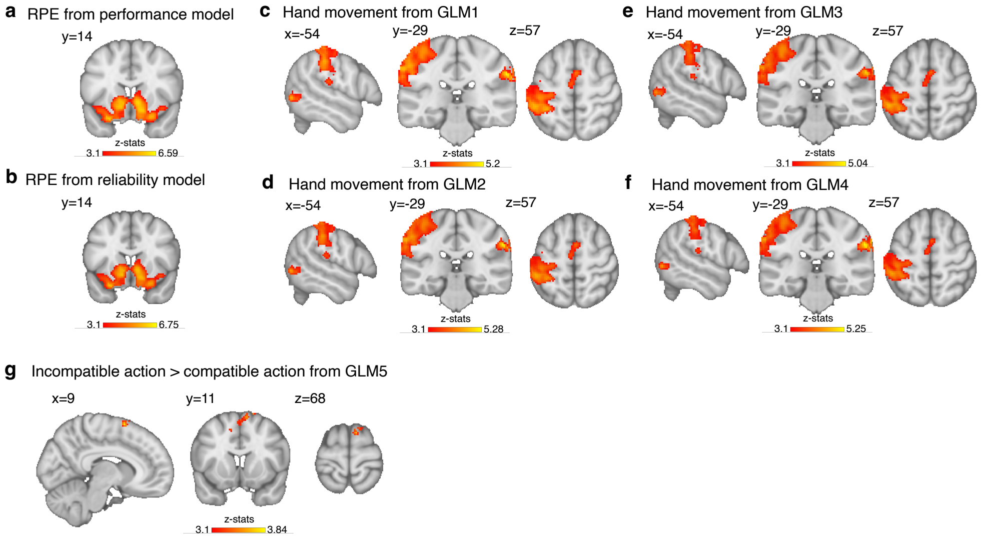


**Supplementary Fig. 14**: Neural correlates of reward prediction error (RPE), motor control and stimulus response incompatibility. (a and b) RPE signals from both models correlated with the ventral striatum. (c to f) The motor control information extracted from the recorded hand videos identified left primary motor cortex and adjacent regions. GLM1 and 3 are the GLMs utilizing variables extracted from the performance-based arbitration model, GLM2 and 4 are the GLMs using variables from the reliability-based arbitration model. All the results were cluster-corrected $p < 0.05$ with the cluster defining threshold $z = 3.1$. (G) The analysis utilizing GLM5 revealed that the dorsal premotor area (PMd) and preSMA exhibit greater activation when the affordance of an object and the selected response are incompatible, compared to when an affordance-compatible action is chosen (uncorrected, $p < 0.001$). See Methods for the details on GLMs.

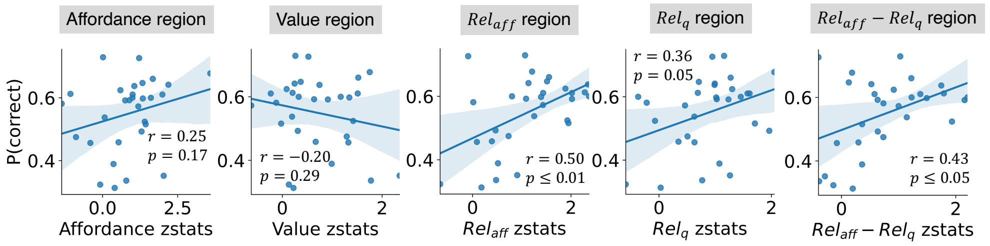


**Supplementary Fig. 15**: Correlations between the strength of the neural representation of cognitive variables and the tendency to select the most rewarding actions. The plots relate to those in Fig. 5 with regions identified using the GLMs based on the reliability-based arbitration model, and the variables of interest are from the reliability-based arbitration model. Each dot represents an individual participant.

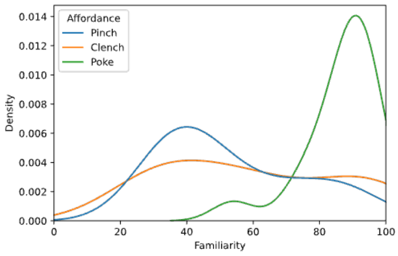


**Supplementary Fig. 16**: Distributions of familiarity scores by the type of affordances of stimuli.

| Behavior study (n=19) | Coef. | Std. Err. | z | P>\|z\| |
|---|---|---|---|---|
| **Intercept** | 6.960 | 0.056 | 124.404 | 0.000 |
| **Choosing pinch** | -0.006 | 0.018 | -0.327 | 0.744 |
| **Choosing clench** | -0.003 | 0.010 | -0.260 | 0.795 |
| **Choosing affordance** | -0.026 | 0.010 | -2.664 | 0.008 |
| **Choosing correct** | -0.054 | 0.017 | -3.143 | 0.002 |
| **Trial index** | -0.006 | 0.001 | -5.970 | 0.000 |
| **In congruent condition** | 0.006 | 0.008 | 0.689 | 0.491 |
| **In high condition** | -0.052 | 0.010 | -5.289 | 0.000 |

| fMRI study (n=30) | Coef. | Std. Err. | z | P>\|z\| |
|---|---|---|---|---|
| **Intercept** | 6.915 | 0.051 | 135.278 | 0.000 |
| **Choosing pinch** | -0.006 | 0.010 | -0.593 | 0.553 |
| **Choosing clench** | -0.018 | 0.015 | -1.174 | 0.241 |
| **Choosing affordance** | -0.029 | 0.010 | -3.016 | 0.003 |
| **Choosing correct** | -0.058 | 0.014 | -4.214 | 0.000 |
| **Trial index** | -0.003 | 0.001 | -2.215 | 0.027 |
| **In congruent condition** | -0.004 | 0.009 | -0.471 | 0.638 |
| **In high condition** | -0.075 | 0.010 | -7.362 | 0.000 |

**Supplementary table 1**: Fixed-effect coefficients of the mixed-effect general linear models on the reaction time. The trial-by-trial reaction time, which is the dependent variable, was logarithmically transformed.

| Behavior study (n=19) | | | |
|---|---|---|---|
| **Initial response** | **Pinch object** | **Clench object** | **Poke object** |
| **Pinch** | 102 (**23**) | 79 (0) | 56 (-23) |
| **Clench** | 93 (-15) | 138 (**30**) | 93 (-15) |
| **Poke** | 109 (-8) | 87 (-30) | 155 (**38**) |

| fMRI study (n=30) | | | |
|---|---|---|---|
| **Initial response** | **Pinch object** | **Clench object** | **Poke object** |
| **Pinch** | 98 (**28.33**) | 62 (-7.67) | 49 (-20.67) |
| **Clench** | 61 (-19.67) | 109 (**28.33**) | 72 (-8.33) |
| **Poke** | 81 (-8.67) | 69 (-20.67) | 119 (**29.33**) |

**Supplementary table 2**: Initial responses sorted by the affordance of the object. The numbers indicate the sum across all initial trials where a particular action was selected as the initial response to the specific object type. The numbers in the parentheses represent the deviation of the number in the cell from the expected number of choosing a particular action type, regardless of the object type which was calculated as the average across columns.

| Behavior study (n=19) | Coef. | Std. Err. | z | P>\|z\| |
|---|---|---|---|---|
| **Intercept** | 0.116 | 0.033 | 3.474 | 0.001 |
| **Number of rewarded trials** | -0.010 | 0.003 | -3.160 | 0.002 |

| fMRI study (n=30) | Coef. | Std. Err. | z | P>\|z\| |
|---|---|---|---|---|
| **Intercept** | 0.113 | 0.035 | 3.216 | 0.001 |
| **Number of rewarded trials** | -0.009 | 0.003 | -2.968 | 0.003 |

**Supplementary table 3**: Fixed-effect coefficients from mixed-effect general linear models on the difference in the frequency of choosing the most-rewarding action between congruent and incongruent conditions. The tables are from the GLMs where the number of rewarded trials previously encountered is the regressor.

| Behavior-generative model \ Identified model | RL | Bayesian | Prior | Bias | Fixed | Cost | Conflict | Rel | Perf |
|---|---|---|---|---|---|---|---|---|---|
| RL | **0.24** | 0 | 0.11 | 0.06 | 0.03 | 0 | 0.03 | 0 | 0 |
| Bayesian | 0.01 | **0.92** | 0 | 0 | 0 | 0 | 0 | 0 | 0 |
| Prior | 0.13 | 0.03 | **0.58** | 0.26 | 0.09 | 0 | 0 | 0.12 | 0 |
| Bias | 0.17 | 0.03 | 0 | **0.36** | 0.03 | 0.13 | 0 | 0 | 0.02 |
| Fixed | 0.11 | 0.02 | 0.05 | 0.06 | **0.43** | 0.13 | 0.31 | 0.03 | 0.04 |
| Cost | 0.11 | 0 | 0.11 | 0.11 | 0.29 | **0.75** | 0.26 | 0.03 | 0.04 |
| Conflict | 0.25 | 0 | 0.05 | 0.02 | 0.03 | 0 | **0.03** | 0.03 | 0 |
| Rel | 0.08 | 0 | 0.05 | 0.06 | 0.06 | 0 | 0.29 | **0.79** | 0 |
| Perf | 0.04 | 0 | 0.05 | 0.06 | 0.06 | 0 | 0.09 | 0 | **0.89** |

**Supplementary table 4**: Probability of the identified model being the actual model that generated the behavior.

Behavior Simulation (n=19x10)

| Fixed Arb. | Coef. | Std. Err. | z | P>lzl |
|---|---|---|---|---|
| Intercept | 0.068 | 0.008 | 8.401 | 0.000 |
| Trial index | -0.000 | 0.001 | -0.301 | 0.763 |

| Cost | Coef. | Std. Err. | z | P>lzl |
|---|---|---|---|---|
| Intercept | 0.047 | 0.007 | 6.483 | 0.000 |
| Trial index | 0.005 | 0.001 | 8.027 | 0.000 |

| Conf. Arb. | Coef. | Std. Err. | z | P>lzl |
|---|---|---|---|---|
| Intercept | -0.012 | 0.007 | -1.750 | 0.080 |
| Trial index | -0.001 | 0.001 | -1.209 | 0.226 |

| Rel. Arb. | Coef. | Std. Err. | z | P>lzl |
|---|---|---|---|---|
| Intercept | 0.071 | 0.009 | 8.151 | 0.000 |
| Trial index | 0.001 | 0.001 | 1.016 | 0.310 |

| Perf. Arb. | Coef. | Std. Err. | z | P>lzl |
|---|---|---|---|---|
| Intercept | 0.109 | 0.010 | 11.015 | 0.000 |
| Trial index | -0.001 | 0.001 | -2.258 | 0.024 |

fMRI Simulation (n=30x10)

| Fixed Arb. | Coef. | Std. Err. | z | P>lzl |
|---|---|---|---|---|
| Intercept | 0.042 | 0.009 | 4.748 | 0.000 |
| Trial index | 0.001 | 0.001 | 1.556 | 0.120 |

| Cost | Coef. | Std. Err. | z | P>lzl |
|---|---|---|---|---|
| Intercept | 0.044 | 0.009 | 5.134 | 0.000 |
| Trial index | 0.003 | 0.001 | 4.603 | 0.000 |

| Conf. Arb. | Coef. | Std. Err. | z | P>lzl |
|---|---|---|---|---|
| Intercept | 0.004 | 0.008 | 0.497 | 0.619 |
| Trial index | -0.001 | 0.001 | -1.214 | 0.225 |

| Rel. Arb. | Coef. | Std. Err. | z | P>lzl |
|---|---|---|---|---|
| Intercept | 0.089 | 0.009 | 9.413 | 0.000 |
| Trial index | -0.001 | 0.001 | -0.943 | 0.346 |

| Perf. Arb. | Coef. | Std. Err. | z | P>lzl |
|---|---|---|---|---|
| Intercept | 0.094 | 0.009 | 10.184 | 0.000 |
| Trial index | -0.001 | 0.001 | -1.500 | 0.134 |

**Supplementary table 5**: Results of mixed-effect general linear models on the simulated behaviors. Fixed-effect coefficients obtained from the mixed-effect GLM on the difference in the choice accuracy between congruent and incongruent conditions are reported, with the trial index being the regressor.

**Behavior Simulation (n=19x10)**

| Model | | Coef. | Std. Err. | z | P>\|z\| |
|---|---|---|---|---|---|
| Fixed Arb. | Intercept | 0.056 | 0.008 | 6.691 | 0.000 |
| | Number of rewarded trials | -0.001 | 0.001 | -0.908 | 0.364 |
| Cost | Intercept | 0.086 | 0.009 | 10.144 | 0.000 |
| | Number of rewarded trials | -0.002 | 0.001 | -2.351 | 0.019 |
| Conf. Arb. | Intercept | -0.025 | 0.007 | -3.513 | 0.000 |
| | Number of rewarded trials | 0.003 | 0.001 | 3.775 | 0.000 |
| Rel. Arb. | Intercept | 0.075 | 0.008 | 9.996 | 0.000 |
| | Number of rewarded trials | -0.003 | 0.001 | -3.077 | 0.002 |
| Perf. Arb. | Intercept | 0.099 | 0.009 | 11.612 | 0.000 |
| | Number of rewarded trials | -0.007 | 0.001 | -6.644 | 0.000 |

**fMRI Simulation (n=30x10)**

| Model | | Coef. | Std. Err. | z | P>\|z\| |
|---|---|---|---|---|---|
| Fixed Arb. | Intercept | 0.037 | 0.008 | 4.410 | 0.000 |
| | Number of rewarded trials | 0.001 | 0.001 | 0.618 | 0.537 |
| Cost | Intercept | 0.072 | 0.008 | 8.871 | 0.000 |
| | Number of rewarded trials | -0.004 | 0.001 | -4.453 | 0.000 |
| Conf. Arb. | Intercept | -0.005 | 0.007 | -0.786 | 0.432 |
| | Number of rewarded trials | -0.000 | 0.001 | -0.054 | 0.957 |
| Rel. Arb. | Intercept | 0.078 | 0.008 | 9.510 | 0.000 |
| | Number of rewarded trials | -0.004 | 0.001 | -4.280 | 0.000 |
| Perf. Arb. | Intercept | 0.097 | 0.008 | 11.536 | 0.000 |
| | Number of rewarded trials | -0.008 | 0.001 | -8.814 | 0.000 |

**Supplementary table 6**: Results of mixed-effect general linear models on the simulated behaviors. Fixed-effect coefficients obtained from the mixed-effect GLM on the difference in the choice accuracy between congruent and incongruent conditions are reported, with the number of rewarded trials previously encountered being the regressor.

**Behavior Simulation (n=19x10)**

| Model | | Coef. | Std. Err. | z | P>\|z\| |
|---|---|---|---|---|---|
| Fixed Arb. | Intercept | -0.024 | 0.007 | -3.263 | 0.001 |
| | Number of rewarded trials | -0.000 | 0.002 | -0.209 | 0.834 |
| Cost | Intercept | -0.051 | 0.007 | -7.192 | <0.001 |
| | Number of rewarded trials | 0.003 | 0.002 | 2.243 | 0.025 |
| Conf. Arb. | Intercept | -0.072 | 0.009 | -8.335 | <0.001 |
| | Number of rewarded trials | 0.006 | 0.001 | 5.557 | <0.001 |
| Rel. Arb. | Intercept | -0.022 | 0.007 | -2.967 | 0.003 |
| | Number of rewarded trials | 0.003 | 0.001 | 1.820 | 0.069 |
| Perf. Arb. | Intercept | -0.038 | 0.008 | -4.762 | <0.001 |
| | Number of rewarded trials | 0.002 | 0.002 | 1.106 | 0.269 |

**fMRI Simulation (n=30x10)**

| Model | | Coef. | Std. Err. | z | P>\|z\| |
|---|---|---|---|---|---|
| Fixed Arb. | Intercept | -0.019 | 0.008 | -2.402 | 0.016 |
| | Number of rewarded trials | 0.003 | 0.001 | 2.248 | 0.025 |
| Cost | Intercept | -0.011 | 0.008 | -1.257 | 0.209 |
| | Number of rewarded trials | -0.000 | 0.001 | -0.209 | 0.834 |
| Conf. Arb. | Intercept | -0.036 | 0.010 | -3.647 | <0.001 |
| | Number of rewarded trials | 0.005 | 0.002 | 3.282 | 0.001 |
| Rel. Arb. | Intercept | 0.001 | 0.008 | 0.109 | 0.913 |
| | Number of rewarded trials | 0.007 | 0.002 | 4.694 | <0.001 |
| Perf. Arb. | Intercept | -0.013 | 0.008 | -1.599 | 0.110 |
| | Number of rewarded trials | 0.003 | 0.002 | 1.672 | 0.094 |

**Supplementary table 7**: Results of mixed-effect general linear models on the simulated behaviors shown in the Supplementary Fig.11. Fixed-effect coefficients obtained from the mixed-effect GLM on the difference in the choice accuracy between high and low conditions are reported, with the number of rewarded trials previously encountered being the regressor.
